# Protein fitness landscapes are simpler under evolutionary distributions

**DOI:** 10.64898/2026.07.08.737351

**Authors:** Darin Tsui, Kunal Talreja, Amirali Aghazadeh

## Abstract

High-order epistasis is often taken as evidence that protein fitness depends on complex interactions among many mutations. The classical Walsh–Hadamard transform quantifies these interactions by weighting all genotypes equally, yet many questions in protein evolution and engineering concern fitness variations among evolutionarily plausible variants. Here we show that weighting genotypes by site-specific amino acid frequencies inferred from antibody and protein language models yields sparser, lower-order representations of protein fitness. Across nine experimental landscapes, the resulting Walsh basis explains fitness variation among evolutionarily plausible variants with, on average, 21% fewer terms overall and 37% fewer terms involving three or more sites than the uniform basis. Sparse regression in the evolutionary basis further explains up to nine percentage points more fitness variation. In three antibody landscapes, mutations at known epistatic hotspots have interpretable single-site effects in the evolutionary basis, while the uniform basis distributes their contributions across multiple interactions. The results show that the reference distribution shapes the apparent complexity of protein epistasis and that weightings derived from evolutionary distributions can make protein fitness landscapes easier to explain and predict.

## Introduction

Understanding how mutations combine to determine protein function remains a central challenge in biology. The effect of a mutation often depends on the sequence background in which it occurs, a phenomenon known as *epistasis*. These interactions shape evolutionary trajectories, constrain accessible mutational pathways, and limit our ability to predict the effects of mutations from sequence [1–3].

Advances in deep mutational scanning have transformed the study of epistasis by allowing quantitative measurements of protein fitness landscapes across thousands to millions of sequence variants [4–6]. These studies have produced two seemingly contrasting views of protein epistasis. On the one hand, many studies have reported extensive higher-order interactions among mutations, suggesting that protein fitness landscapes are highly nonlinear and combinatorially complex [7–11]. On the other hand, a growing body of work has shown that genotype–phenotype relationships can exhibit weak effective higher-order epistasis, low-dimensional organization, and sparse representations that enable accurate prediction from relatively few measurements [3, 12–15]. In some proteins, models of global epistasis explain much of the measured variation through a nonlinear relationship between an additive trait and the observed phenotype [16]. However, how complex a landscape appears depends on the protein, the representation used, and the region of sequence space under study.

When the goal is to characterize epistasis throughout the fitness landscape, it is natural to weigh each genotype equally. The classical Walsh–Hadamard transform (WHT) does this by forming an orthonormal basis under the uniform measure, separating single-site effects from interactions of increasing order [8]. When relatively few terms explain most of the fitness variation, this representation enables landscape reconstruction from incomplete measurements and provides a natural way to regularize models of sequence–function relationships [15, 17–20].

Many questions about protein epistasis, however, concern a particular region of the landscape. In evolutionary studies, we may ask how mutations interact across the sequence backgrounds that natural proteins could plausibly occupy [21]. In protein design, we may want to rank functional variants by binding affinity or stability [22]. Both questions require resolving fitness differences among variants that occupy only a small part of sequence space, a part that the uniform measure weights no more heavily than the rest. To model variation in this region, one can instead weight genotypes by the site-specific amino acid frequencies of evolutionarily plausible sequences.

Quantitative genetics has long recognized that estimates of epistasis depend on the allele frequencies of the population in which they are measured [23, 24]. For populations in linkage equilibrium, where genotype frequencies factor into site-specific allele frequencies, Stadler and colleagues showed that each term of the uniform Walsh–Hadamard transform contributes only to the term involving the same sites and to lower-order terms involving subsets of those sites [25]. Reweighting is therefore expected to move weight toward lower orders. Because a single interaction can contribute to several terms, and contributions from different interactions can reinforce or cancel, the same reweighting can make the representation either sparser or denser. It remains an open question whether evolutionary weighting simplifies the representation of experimentally measured protein fitness landscapes.

Here we show that evolutionary weighting yields representations of protein fitness landscapes that are both lower-order and sparser. We used the antibody language model AntiBERTy [26] and the protein language model ESM2 [28] to identify evolutionarily plausible variants and estimated site-specific mutant frequencies among them. The product of these frequencies defines an evolutionary distribution over sequence space and a corresponding *p*-biased Walsh–Hadamard transform, which we call the *evolutionary Walsh–Hadamard transform* (eWHT). The classical WHT is the special case in which every site has *p*_*i*_ = 0.5 (Fig. 1). We compared the two transforms on the same language model-selected variants in nine experimentally measured fitness landscapes. To explain 95% of fitness variation, eWHT required fewer terms than WHT in eight of the nine landscapes. Across landscapes, eWHT required 21% fewer terms overall and 37% fewer terms involving three or more sites. We show the important consequences of this result: with the same number of measurements, sparse regression in the evolutionary basis explains up to nine percentage points more fitness variation. Randomizing the sequence–fitness relationship while holding the evolutionary weights and evaluation variants fixed, or shuffling the evolutionary weights while holding the fitness landscape and evaluation variants fixed, weakened or reversed the gain. These findings suggest that part of the apparent complexity of protein epistasis reflects the reference distribution against which it is measured, and that fitness among evolutionarily plausible variants may now be learnable from fewer experiments.

**Figure 1:**
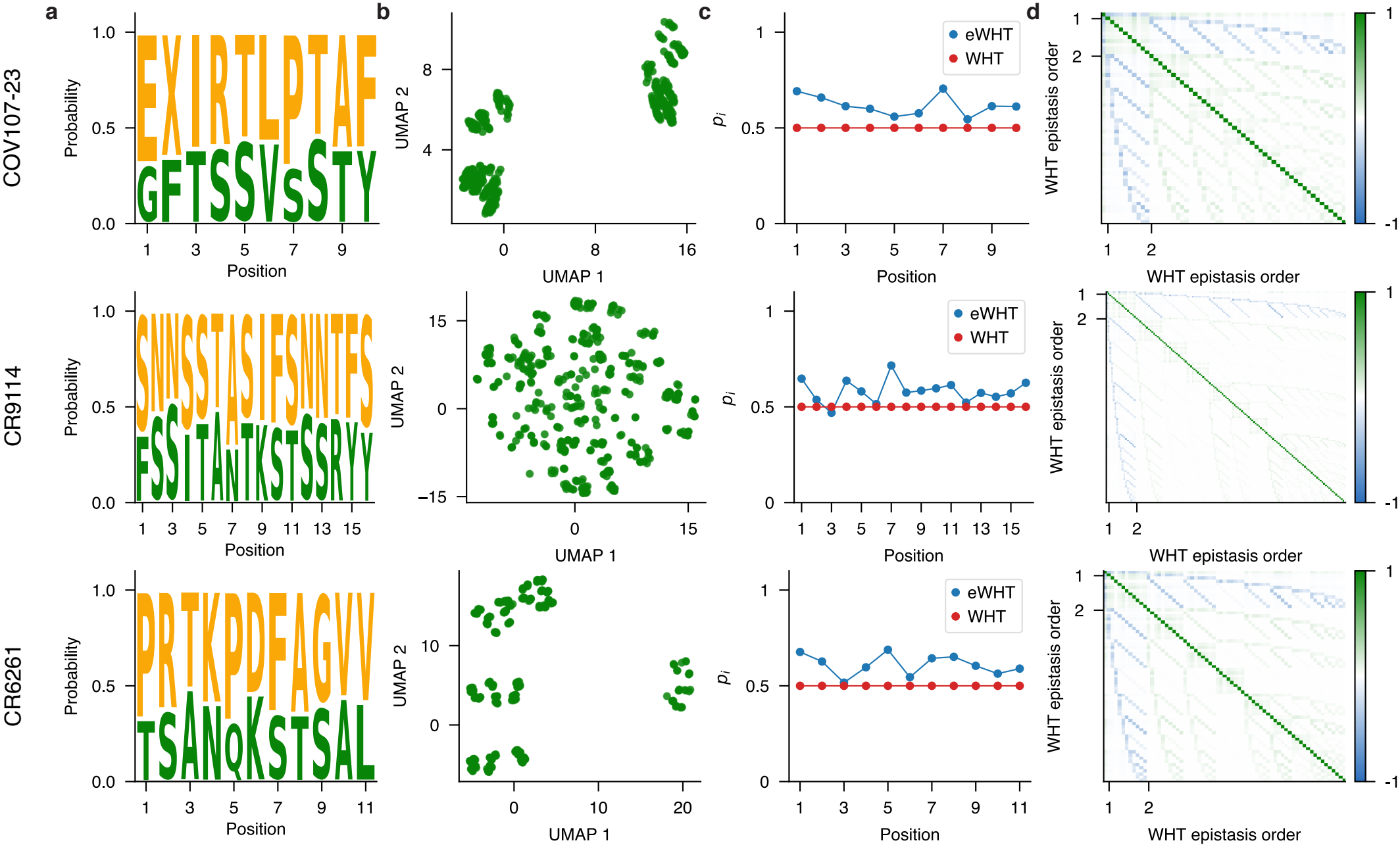
Evolutionarily plausible variants occupy a non-uniform region of sequence space, motivating an evolutionary Walsh–Hadamard transform (eWHT). Variants were selected using the antibody language model AntiBERTy [26] for the SARS-CoV-2-specific antibody COV107-23 (top) [10] and the broadly neutralizing influenza antibodies CR9114 (middle) and CR6261 (bottom) [27]. **a** Wild-type (green) and mutant (orange) amino acid frequencies at each site among the selected variants. Many sites strongly favor one residue. **b** UMAP projections of AntiBERTy embeddings for the selected variants. **c** Mutant frequency *p*_*i*_ at each site *i*, defined as the fraction of selected variants carrying the mutant residue. Across all nine landscapes, *p*_*i*_ ranges from 0.13 to 0.76 (see Supplementary Information). The classical Walsh–Hadamard transform (WHT) implicitly assumes *p*_*i*_ = 0.5 at every site. **d** Gram matrix of the classical WHT basis functions through second order, evaluated under the product measure defined by the frequencies in **c**. Nonzero off-diagonal entries show that the classical basis is no longer orthogonal once genotypes are weighted by these frequencies, so its terms no longer separate cleanly. The eWHT basis is orthonormal under this measure by construction, and it recovers the classical WHT when *p*_*i*_ = 0.5 at every site.

## Results

### An Evolutionary Basis for Protein Fitness Landscapes

To define an evolutionary reference distribution for each fitness landscape, we scored all variants with the antibody language model AntiBERTy [26] or the protein language model ESM2-650M [28] and selected a subset, which we refer to as *evolutionarily plausible* variants (see Methods). Fig. 1 illustrates this procedure for three antibody landscapes scored with AntiBERTy. Among the selected variants, the frequencies of wild-type and mutant residues varied substantially across sites (Fig. 1a), and UMAP projections of their AntiBERTy embeddings formed distinct clusters (Fig. 1b) [29].

We summarized each selected set by the mutant frequency *p*_*i*_ at each site *i*, defined as the fraction of selected variants carrying the mutant residue, and collected these into the evolutionary probability vector ***p*** = [*p*_1_, *p*_2_, …, *p*_*L*_]. These frequencies departed markedly from the value *p*_*i*_ = 0.5 implicit in the classical Walsh–Hadamard transform (WHT) (Fig. 1c). Let *f* (**x**) denote the fitness of genotype **x** ∈ {−1, 1}^*L*^, where *x*_*i*_ = 1 and *x*_*i*_ = −1 denote the wild-type and mutant states at site *i*. The frequencies define a product measure over the full landscape,

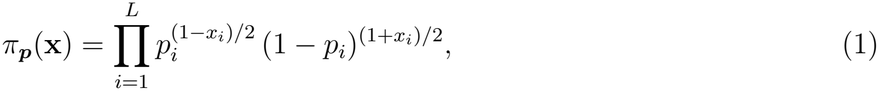

which assigns each genotype the probability of drawing its residues independently at each site. Under this measure, the classical Walsh–Hadamard basis is no longer orthogonal. The corresponding Gram matrix has nonzero off-diagonal entries (Fig. 1d), so its coefficients no longer partition fitness variation into separate single-site effects and interactions. We therefore represented fitness in the basis that is orthonormal under *π*_***p***_, which we call the *evolutionary Walsh–Hadamard transform* (eWHT). For 0 *< p*_*i*_ *<* 1 at every site, the eWHT expresses fitness as:

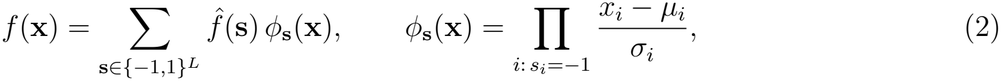

where *µ*_*i*_ = 1 − 2*p*_*i*_ and 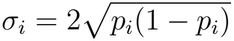 are the mean and standard deviation of *x*_*i*_ under *π*_***p***_. Each coefficient 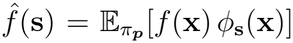 is the projection of fitness onto *ϕ*_**s**_, and the number of sites with *s*_*i*_ = −1 defines the order of the corresponding term. When *p*_*i*_ = 0.5 at every site, *µ*_*i*_ = 0 and *σ*_*i*_ = 1, and Equation (2) reduces to the classical Walsh–Hadamard transform. Each eWHT term thus retains the interpretation of a single-site effect or an interaction among a specific set of sites, but measures these effects relative to the evolutionary distribution rather than the uniform distribution. The matrix formulation and an *O*(*N* log *N*) algorithm for computing the transform, where *N* = 2^*L*^, are given in Methods.

### Evolutionary Distributions Simplify Protein Fitness Landscapes

To determine how evolutionary weighting affects the representation of protein fitness landscapes, we analyzed nine combinatorially complete deep mutational scanning datasets, for which both the eWHT and WHT can be computed exactly. These landscapes comprise the folding stability of the SARS-CoV-2-specific antibody COV107-23 [10], the binding affinities of the influenza broadly neutralizing antibody CR9114 to hemagglutinin H1 and H3 (CR9114-H1 and CR9114-H3) and of CR6261 to hemagglutinins H1 and H9 (CR6261-H1 and CR6261-H9) [27], TagBFP fluorescence [3], His3p growth rate [21], *β*-lactamase cefotaxime resistance [1], and angiotensin-converting enzyme 2 (ACE2) affinity to the Hu-1 receptor-binding domain [9]. Fig. 2 shows three representative landscapes: COV107-23, CR9114-H1, and CR6261-H1. Results for the remaining landscapes are given in the Supplementary Information.

**Figure 2:**
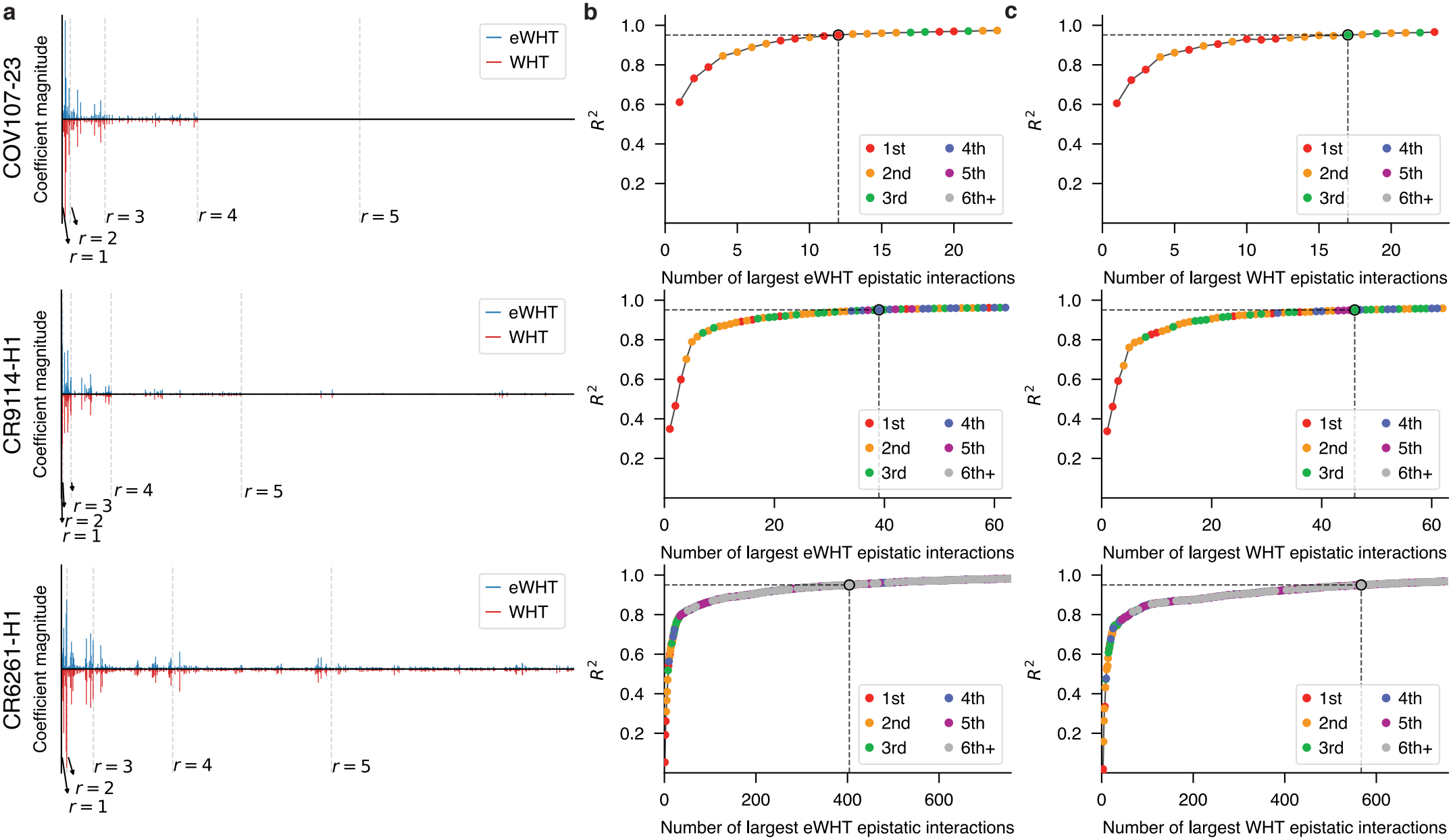
Evolutionary weighting yields sparser, lower-order representations of protein fitness landscapes. The evolutionary Walsh–Hadamard transform (eWHT; upper plots, blue) is compared with the classical Walsh–Hadamard transform (WHT; lower plots, red) for three representative landscapes: the SARS-CoV-2-specific antibody COV107-23 (top) and the influenza broadly neutralizing antibodies CR9114 (middle) and CR6261 (bottom), both binding hemagglutinin subtype H1. **a** Magnitudes of eWHT and WHT coefficients, grouped by interaction order. Dashed vertical lines separate orders one through five. **b** Fraction of fitness variation explained (*R*^2^) among evolutionarily plausible variants as eWHT terms are added in decreasing order of coefficient magnitude. Colors indicate interaction order, and dashed lines mark the smallest number of terms that reaches *R*^2^ ≥ 0.95. **c** The same analysis for WHT terms on the same variants. In each landscape shown, eWHT reaches the threshold with fewer terms than WHT and relies less on interactions among three or more sites.

For each landscape, we defined the evolutionary measure ***p*** using AntiBERTy [26] or ESM2-650M [28], language models that score the plausibility of each variant under the distribution of natural protein sequences. For the five antibody landscapes (COV107-23, CR9114-H1, CR6261-H1, CR9114-H3, and CR6261-H9), we used AntiBERTy and scored each variant by its pseudo-perplexity [30, 31]. For the four protein landscapes (TagBFP, His3p, Hu-1, and *β*-lactamase), we used ESM2-650M and scored each variant by the sum of the log-likelihood ratios of its mutations relative to the wild-type sequence [32]. Following a convention similar to that of [33], we designated half of the variants as evolutionarily plausible and set *p*_*i*_ to the fraction of these variants carrying the mutant residue at site *i* (Fig. 1c).

We then compared how compactly the two transforms represent measured fitness on two sets of variants: *(i)* the evolutionarily plausible variants defined as above and *(ii)* experimentally high-fitness variants, defined as those with measured fitness at or above the median [33]. For the evolutionarily plausible variants, we added terms in decreasing order of coefficient magnitude and computed the fraction of fitness variation explained (*R*^2^) after each addition (Fig. 2b,c; Methods). eWHT reached *R*^2^ = 0.95 with fewer terms than WHT in eight of the nine landscapes. Listed in decreasing order of the absolute Spearman correlation |*ρ*| between the language model score and measured fitness, eWHT required 29% fewer terms in CR6261-H1 (|*ρ*| = 0.62), 67% in *β*-lactamase (|*ρ*| = 0.61), 23% in CR6261-H9 (|*ρ*| = 0.53), 29% in COV107-23 (|*ρ*| = 0.44), 15% in CR9114-H1 (|*ρ*| = 0.34), 22% in His3p (|*ρ*| = 0.31), 3% in CR9114-H3 (|*ρ*| = 0.26), and 7% in Hu-1 (|*ρ*| = 0.14). TagBFP (|*ρ*| = 0.12), the landscape with the weakest correlation, was the only exception, requiring 4% more terms. The three smallest reductions occurred in the three landscapes where the score correlated least with fitness, suggesting that eWHT’s success in sparsifying representations depends on how well the evolutionary distribution aligns with fitness. Averaged over the six landscapes in which the score correlated with fitness at least moderately, which we define as Spearman |*ρ*| *>* 0.3, eWHT required 31% fewer terms overall and 48% fewer terms involving three or more sites. Across all nine landscapes, eWHT required 21% fewer terms overall and 37% fewer terms involving three or more sites.

Among high-fitness variants, fitness varies over a narrow range, making *R*^2^ sensitive to small absolute errors, so we assessed this set by mean squared error (MSE) instead (Fig. 3). eWHT reached an MSE of 0.05 with fewer terms than WHT in five of the nine landscapes, with two ties, and required 13% fewer terms overall on average. Only CR6261-H1, CR9114-H3, and CR6261-H9 required third-order or higher terms under WHT at this threshold, and in these three landscapes eWHT required 30% fewer such terms on average. Under the evolutionary distribution, measured fitness is captured by fewer terms, with the largest reduction among high-order interactions. This holds for both evolutionarily plausible variants and the variants that perform best experimentally.

**Figure 3:**
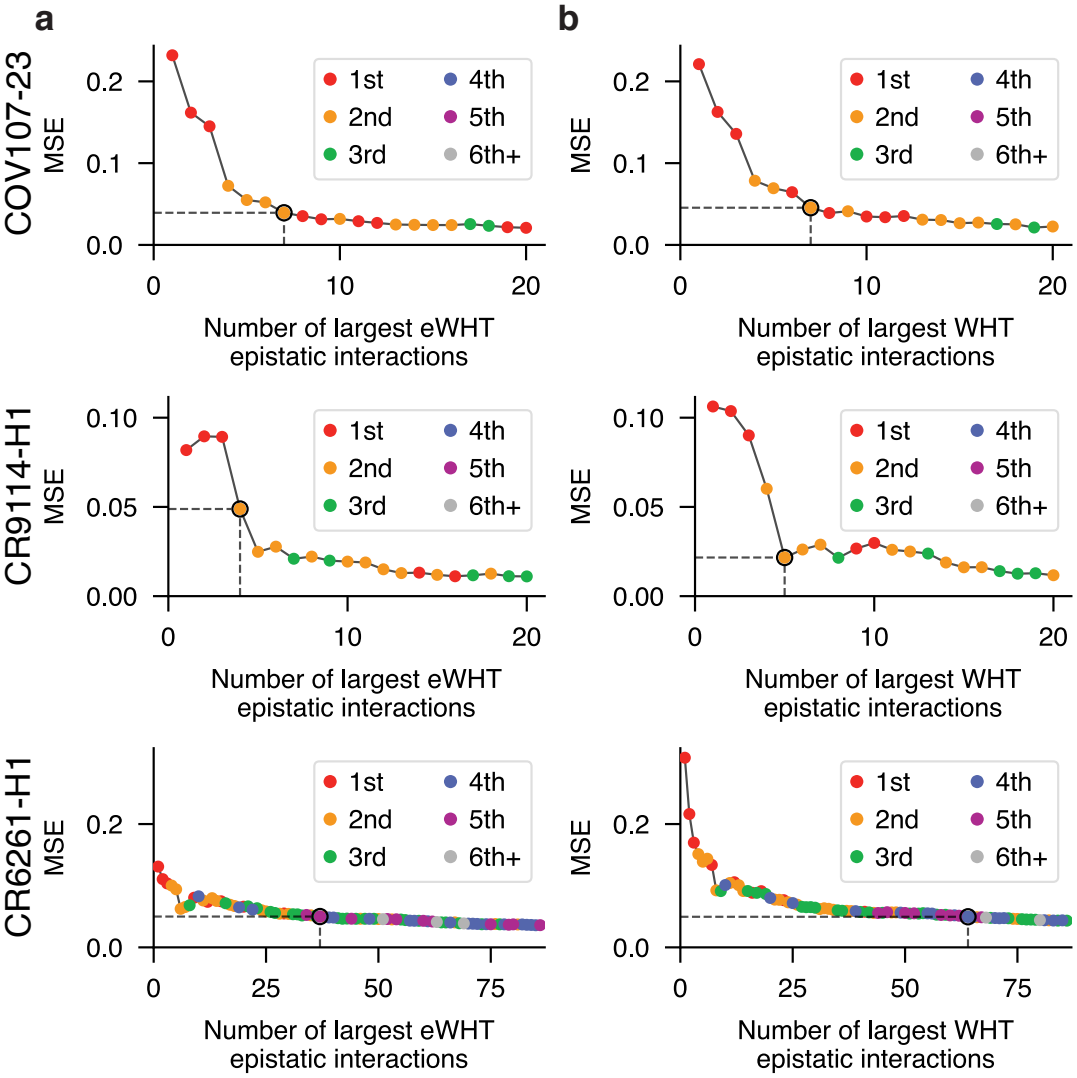
Evolutionary weighting yields sparser representations among experimentally high-fitness variants. Mean squared error (MSE) among variants with measured fitness at or above the median, as terms are added in decreasing order of coefficient magnitude to the **a** eWHT and **b** classical WHT expansions, for COV107-23 (top), CR9114-H1 (middle), and CR6261-H1 (bottom). Colors indicate interaction order. Black dots and dashed lines mark the smallest number of terms that reaches MSE ≤ 0.05.

To test whether the sparsity gain from eWHT depends on how fitness values are assigned to genotypes and on the evolutionary measure, we repeated the compactness analysis on five synthetic controls for each experimental landscape: three that altered the genotype–fitness map while holding the evolutionary measure fixed and two that altered the evolutionary measure while holding the genotype– fitness map fixed (see Methods and Supplementary Information). The largest average reductions under the controls that altered the genotype–fitness map were 7% in the total number of terms and 13% in terms involving three or more sites, compared with 21% and 37% on the experimental landscapes. Some controls reversed this advantage, with eWHT requiring up to 273% more total terms and up to 266% more terms involving three or more sites than WHT. The largest average reductions under the controls that altered the evolutionary measure were 12% and 25%. As with the controls that altered the genotype–fitness map, one control also reversed this advantage, with eWHT requiring *>* 80× as many total terms and *>* 250× as many terms involving three or more sites as WHT. Since each control altered either the genotype–fitness map or the evolutionary measure while holding the other and the evaluation variants fixed, these results show that the sparsity gain depends on both. The smaller gains under the controls suggest that the evolutionary measure captures sequence preferences relevant to measured fitness.

### Sparser Representations Improve Fitness Prediction from Limited Measurements

A sparser representation should be easier to learn from incomplete data. When only a small number of terms carry most of the fitness variation, sparse recovery methods can estimate those terms from far fewer measurements than the size of the landscape [15]. If the evolutionary basis concentrates fitness variation into fewer terms than the classical basis, models fitted in it should therefore predict fitness more accurately from the same number of experiments.

To test this, we fitted sparse models in both bases using the least absolute shrinkage and selection operator (LASSO) [34]. For each landscape except *β*-lactamase, which we omitted because its sequence space contains only 32 variants, we drew training sets of increasing size at random from the evolutionarily plausible variants, selected the regularization strength by cross-validation within each training set and separately for each basis, and evaluated accuracy (*R*^2^) on held-out evolutionarily plausible variants not used for training (see Methods). Because the two bases span the same space of fitness functions and are fitted to the same measurements, any difference in accuracy reflects how compactly each basis represents the landscape, not differences in model capacity or tuning.

Across all eight landscapes, models fitted in the evolutionary basis generally matched or exceeded the accuracy of those fitted in the classical basis (Fig. 4a). With 65 measurements in COV107-23 and 300 in CR9114-H1 and CR6261-H1, the evolutionary basis reached *R*^2^ values of 0.80, 0.82, and 0.83, compared with 0.75, 0.77, and 0.74 in the classical basis (Fig. 4b,c). Equivalently, the evolutionary basis attained a given accuracy from fewer measurements. By concentrating fitness variation into fewer terms, eWHT reduces the experimental effort needed to model fitness among evolutionarily plausible variants.

**Figure 4:**
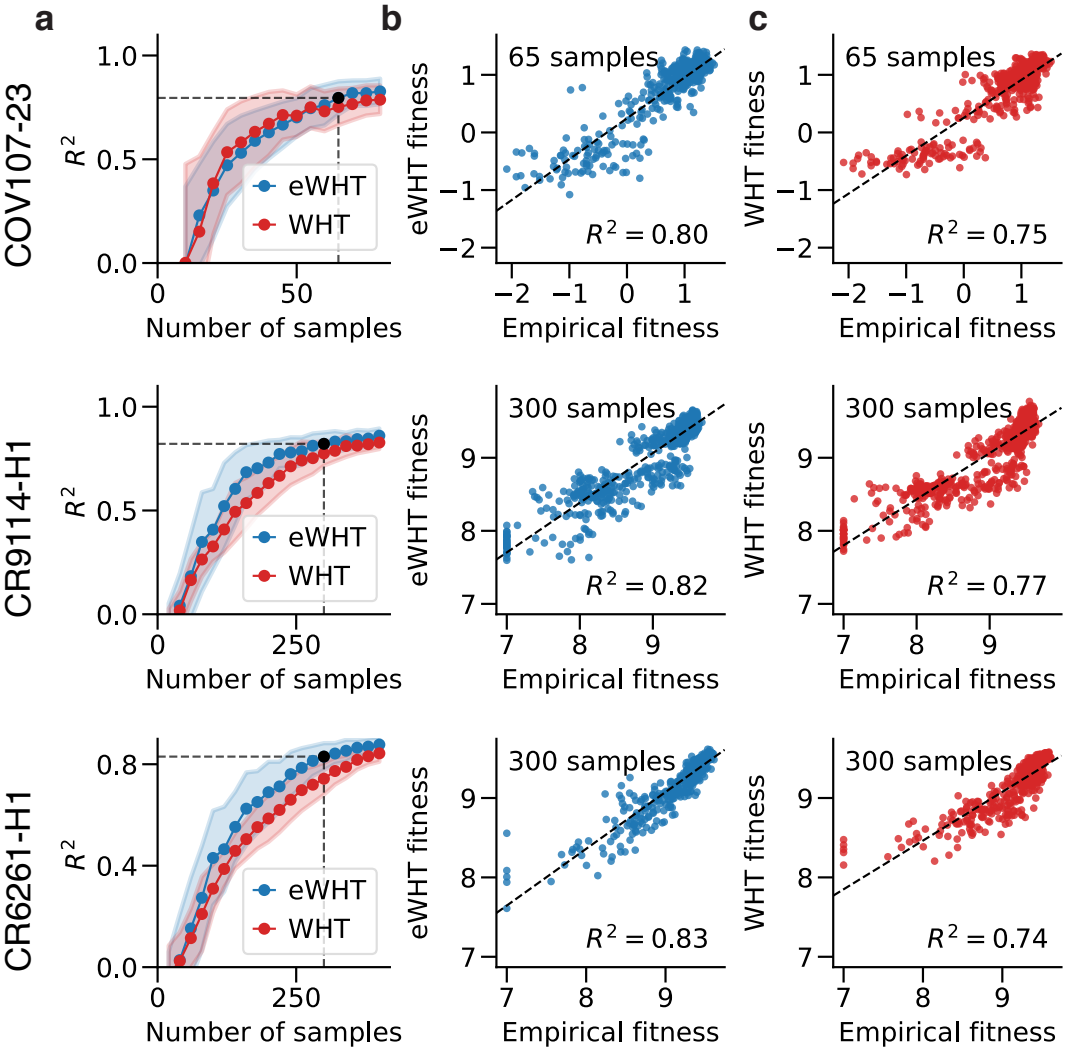
Evolutionary weighting improves reconstruction of protein fitness landscapes from limited experimental data. **a** Accuracy (*R*^2^) on held-out evolutionarily plausible variants of sparse models fitted in the eWHT (blue) and classical WHT (red) bases, as a function of the number of measured training variants. Points show the mean and shaded regions show ± standard deviation across 50 random training draws. In each landscape shown, eWHT generally matches or exceeds WHT in *R*^2^, with the largest improvement in CR6261-H1. **b** Predicted versus measured fitness for models fitted in the eWHT basis. **c** The same comparison for models fitted in the WHT basis. Predictions are shown at the smallest training size at which eWHT reaches 95% of its average *R*^2^ at the largest training size tested (65 variants for COV107-23 and 300 for CR9114-H1 and CR6261-H1).

### Evolutionary Weighting Consolidates Epistasis onto Fewer, Interpretable Terms

To examine how the evolutionary measure changes the interpretation of epistasis, we compared the terms retained by eWHT and WHT when each representation was truncated at *R*^2^ = 0.95 in the three representative landscapes (Fig. 2b,c). At the pairwise level, the two representations largely agreed: many dominant interactions were shared, and the coefficients of shared interactions were strongly correlated (Spearman *ρ* = 1.00, 0.98, and 0.97 in COV107-23, CR9114-H1, and CR6261-H1, respectively; Fig. 5a). eWHT nonetheless yielded a sparser representation among pairwise interactions, retaining 38%, 15%, and 5% fewer interactions than WHT in the respective landscapes.

**Figure 5:**
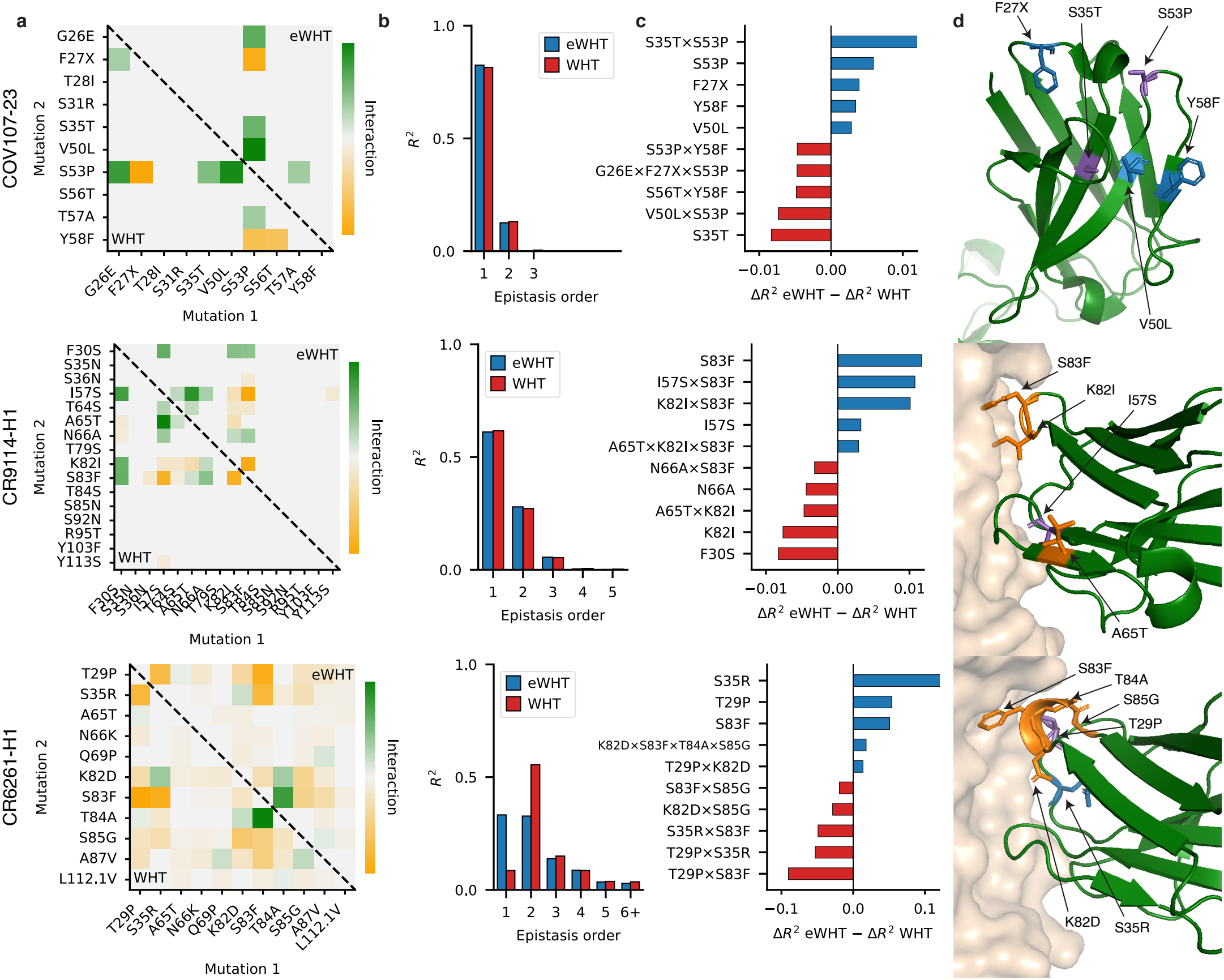
Evolutionary weighting consolidates epistasis onto a few key mutations. Terms retained by eWHT and WHT when each expansion is truncated at *R*^2^ = 0.95 among evolutionarily plausible variants in the COV107-23, CR9114-H1, and CR6261-H1 landscapes (Fig. 2b,c). **a** Pairwise coefficients retained by eWHT (upper triangle) and WHT (lower triangle). The X mutation in COV107-23 denotes a mutation to V, L, or I. eWHT preserves most dominant pairwise interactions while retaining fewer of them overall. **b** *R*^2^ contributed by the retained terms, stratified by interaction order. The contribution of each order is nearly unchanged under eWHT in COV107-23 and CR9114-H1, whereas in CR6261-H1 eWHT shifts contribution from pairwise to first-order terms. **c** The five terms with the largest increase in incremental *R*^2^ under eWHT (positive) and under WHT (negative), shown as 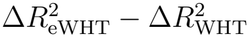. Incremental *R*^2^ is the change in the variance-explained curve of Fig. 2b,c when a term is added. **d** Residues participating in the five largest eWHT gains from **c**, mapped onto each structure and colored by the highest interaction order in which each participates: first (blue), second (purple), and third or higher (orange). Top, COV107-23 (PDB ID 7LKA [35]); middle, CR9114-H1 (PDB ID 4FQI [36]); bottom, CR6261-H1 (PDB ID 3GBN [37]).

To quantify how the two transforms distribute explained variance across interaction orders, we computed the *R*^2^ contributed by each order within the set of terms required to reach *R*^2^ = 0.95 (Fig. 5b). In COV107-23 and CR9114-H1, the contribution of each order under eWHT was nearly unchanged even though eWHT required 29% and 15% fewer terms overall, indicating that it preserved the order structure of these landscapes while consolidating it into fewer interactions. The largest changes occurred in CR6261-H1 and TagBFP, the two landscapes in which WHT assigned the most explained variance to pairwise and higher terms (*R*^2^ = 0.86 and 0.84, respectively, compared with at most 0.67 elsewhere). In CR6261-H1, eWHT moved explained variance from pairwise to first-order terms, reducing the pairwise-and-higher contribution from *R*^2^ = 0.86 to 0.62 and raising the first-order contribution from 0.09 to 0.33. In TagBFP, it reduced the pairwise-and-higher contribution from *R*^2^ = 0.84 to 0.57 and raised the first-order contribution from 0.11 to 0.38 (see Supplementary Information).

To identify which terms each transform emphasizes, we computed the incremental *R*^2^ of each term, defined as the change in the variance-explained curve (Fig. 2b,c) when that term is added, and compared the two representations term by term (Fig. 5c). In COV107-23, the largest eWHT gains were the first-order term of S53P and the interaction S35T×S53P. S53P has been reported as an epistatic hotspot and the most stabilizing single mutation in structure-based stability simulations [10]. Because S53P modulates the fitness effects of nearly all other mutations, WHT distributes its contribution across many pairwise and higher-order terms, including V50L×S53P, G26E×F27X×S53P (where X denotes a mutation to V, L, or I [10]), and S53P×Y58F. The one interaction that eWHT retains pairs S53P on the heavy-chain complementarity-determining region (HCDR) 2 loop with S35T on HCDR1, a site previously characterized as weakly epistatic [10], suggesting that the evolutionary measure isolates the interactions most relevant to the structural stability of this antibody.

The same consolidation was evident in the two influenza antibodies. In CR9114-H1, the largest eWHT gains involved four of the five mutations previously identified as key epistatic sites: I57S, A65T, K82I, and S83F [27]. The fifth, F30S, appeared among the largest WHT gains instead. Because the effect of F30S depends on S83F and I57S [27], whose first-order terms are among the largest eWHT gains, eWHT may absorb part of its contribution into those terms. In CR6261-H1, every term in Fig. 5c involves mutations in CDR1 (T29P and S35R) or framework region (FW) 3 (K82D, S83F, T84A, and S85G), the two regions previously reported to dominate higher-order epistasis in this landscape [27]. The largest eWHT gains were the first-order terms of S35R, T29P, and S83F, mutations that carry the largest third- and fourth-order contributions under WHT, together with a single fourth-order interaction among four consecutive FW3 residues (K82D, S83F, T84A, and S85G). S85G and K82D, absent from the largest eWHT first-order gains, appear under WHT in the pairwise terms S83F×S85G and K82D×S85G, suggesting that eWHT represents these residues jointly through the fourth-order term rather than through separate pairwise ones. Across all three landscapes, evolutionary weighting thus consolidates the many terms that WHT requires into a smaller set centered on a few mutations already known to govern each landscape.

## Discussion

The apparent complexity of a protein fitness landscape depends on which variants the representation is meant to describe and how those variants are weighted when defining epistatic effects. Among evolutionarily plausible variants, the eWHT required 21% fewer terms overall and 37% fewer terms involving three or more sites to explain the same fraction of measured fitness variation as the classical WHT. These reductions are averages of the per-landscape percentage reductions. The change in interaction order varied by landscape. The largest shift toward single-site effects occurred in a landscape with substantially higher-order epistasis under uniform weighting. In landscapes where single-site effects already explained much of the fitness variation, the distribution of explained variance across orders changed less.

Quantitative genetics has long recognized that estimates of epistasis depend on allele frequencies [23, 24]. For a product measure, Stadler and colleagues showed that a term in the uniform expansion can contribute to terms involving the same sites or subsets of those sites [25]. Whether the resulting expansion is sparser depends on how these contributions combine. We found that antibody and protein language model-derived weights yielded sparser representations of measured fitness. The sparsity depends on the degree of alignment between the language model scores and measured fitness. Importantly, randomizing the genotype–fitness map or evolutionary measure weakened or reversed this gain even with the evaluation variants held fixed. These controls show that the gain depends on the fitness map and suggest that the sequence preferences captured by the evolutionary measure are relevant to the measured effects of mutations.

The appropriate measure depends on the problem. Uniform weighting gives every genotype equal importance when modeling fitness across the full landscape. Evolutionary weighting is useful when the objective is to resolve fitness differences among evolutionarily plausible variants, such as differences in stability or binding affinity. In our analysis, we used the site frequencies of antibody and protein language model-selected variants to define the product measure, and we evaluated both representations on the same selected variants. We also found a smaller reduction in sparsity when evaluating the representation on experimentally high-fitness variants. In practice, the sequence distribution used to define the basis, the variants available for training, and the variants on which predictions are needed can differ [38]. In particular, the performance of the evolutionary basis on high-fitness variants far from the sequences favored by the language model remains an open question.

Global epistasis provides another explanation for the high interaction orders inferred from measured phenotypes. A nonlinear relationship between an additive latent trait and the measured phenotype can produce apparent interactions among many sites. Modeling this nonlinearity can reduce the inferred interaction order [16, 21, 39]. Evolutionary weighting changes the reference distribution used to define epistatic effects, while global-epistasis models account for nonlinearities in how mutational effects map to the measured phenotype. An open question is whether accounting for both can further reduce the number or order of terms needed to describe fitness.

Our formulation uses a product measure, which permits independent estimation of site frequencies and an O(*N* log *N*) transform, where *N* is the number of genotypes. A product measure omits dependencies among sites, including those associated with structural contacts and functional constraints [40, 41]. Future work could test whether measures that capture such dependencies yield more compact representations. Extending the transform to *q*-ary alphabets would also allow distinct amino acid substitutions at each site to be represented explicitly [18, 42].

We provide an open-source implementation of the eWHT at https://github.com/amirgroup-codes/ewht, including the fast transform, sparse recovery routines, and utilities for estimating evolutionary measures from antibody and protein language models. At the same experimental budget, sparse regression in the evolutionary basis explained more fitness variation on held-out variants. Together with prior work linking spectral sparsity to recovery from incomplete measurements [15], this result suggests that matching the measure to the sequences of interest can reduce the number of experiments needed to resolve fitness differences among evolutionarily plausible variants.

## Methods

### Evolution-aware Representation of Sequence Space

The evolutionary Walsh–Hadamard transform (eWHT) is an instance of the classical *p*-biased Walsh–Hadamard transform (*p*-WHT), where the bias parameters are estimated from evolutionary sequence distributions (see Section 8.4 of [43]). Unlike the classical Walsh–Hadamard transform, which assumes a uniform probability measure over sequence space, eWHT defines orthonormality with respect to an evolutionary probability measure inferred from antibody and protein language models.

Let 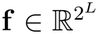 denote the vector of fitness values of all sequences and 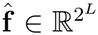 the vector of eWHT coefficients. Given the vector of evolutionary probabilities ***p***, we define the inverse eWHT as:

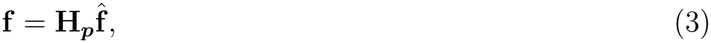

where 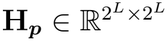 denotes the eWHT basis matrix. Following the classical WHT formulation, **H** can be computed via the Kronecker product:

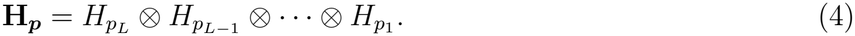

Here, the eWHT Hadamard matrix at position *i*, 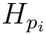, is defined as:

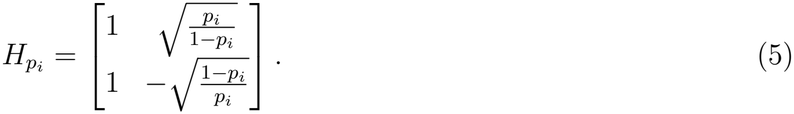

Conversely, given **f**, we define the forward eWHT as:

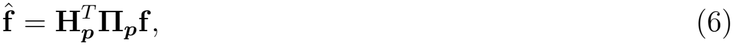

where 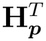 is the transpose of **H**_***p***_ and **Π**_***p***_ = diag (*π*_***p***_(**x**)) is a diagonal expectation weighting matrix representing the joint evolutionary probabilities over sequence space. Because *π*_***p***_ is a product measure, **Π**_***p***_ decomposes via the Kronecker product:

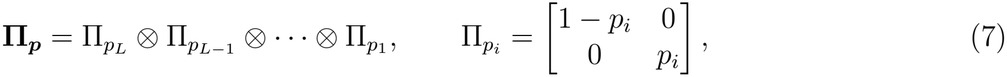

where 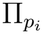 is the single-site weighting matrix, whose entries are the probabilities of the wild-type and mutant states at site *i*.

We can equally represent the forward and inverse eWHT for the sequence **x** ∈ {−1, 1}^*L*^. Given ***p***, the evolutionary probability measure *π*_***p***_(**x**) as defined in Equation (1) specifies the probability of observing sequence **x** under the evolutionary distribution. *π*_***p***_(**x**) allows us to define the corresponding evolutionary inner product as:

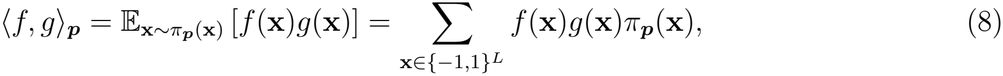

under which the eWHT basis functions are orthonormal. Consequently, every fitness function admits a unique orthonormal decomposition under the evolutionary measure. In particular, the forward eWHT computes the spectral coefficient 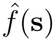, where **s** ∈ {−1, 1}^*L*^, via:

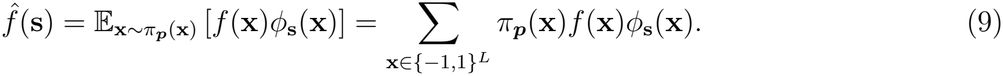

Conversely, the inverse eWHT computes *f* (**x**) via Equation (2). Notably, when *p*_*i*_ = 0.5 for every position, the evolutionary measure reduces to the uniform distribution and eWHT exactly recovers the classical forward and inverse Walsh–Hadamard transforms. Additional properties of the eWHT basis functions, proofs of orthonormality, and uniqueness of the eWHT decomposition are provided in the Supplementary Information.

### Fast Computation of the Evolutionary Walsh–Hadamard Transform

The eWHT coefficients were computed using a fast recursive algorithm that exploits the product structure of the basis functions. Similar to the classical fast Walsh–Hadamard transform, the algorithm admits an implementation with computational complexity *O*(*N* log *N*), where *N* = 2^*L*^. The derivation of the recursion, proofs of correctness, and forward and inverse algorithms are provided in the Supplementary Information.

### Protein Fitness Landscapes

We evaluated eWHT on nine combinatorially complete protein fitness landscapes spanning protein stability, molecular binding, fluorescence, enzyme activity, and cellular growth. These landscapes include the folding stability of COV107-23 [10], the binding affinities of the influenza broadly neutralizing antibodies CR6261 to hemagglutinins H1 (CR6261-H1) and H9 (CR6261-H9) and CR9114 to H1 (CR9114-H1) and H3 (CR9114-H3) [27], TagBFP fluorescence [3], His3p growth rate [21], *β*-lactamase antibiotic resistance [1], and ACE2 affinity to the Hu-1 receptor-binding domain [9]. For the His3p growth rate dataset, we followed the preprocessing steps detailed in [15] to ensure a combinatorially complete dataset. For the remaining datasets, we followed the protocols described in the original publications for experimental measurements and preprocessing. In some datasets, the original publications did not report the fitness of a tiny fraction (typically less than 1% of the landscape) of sequences. For the CR6261-H1, CR6261-H9, CR9114-H1, and CR9114-H3 landscapes, unreported sequences were assigned baseline fitness values of 7.0, 7.0, 6.0, and 6.0, respectively, following the original paper’s convention [27]. For all other datasets, we estimated the fitness of each unreported sequence as the average fitness of sequences that differ only at a subset of the mutated sites.

### Estimating Evolutionary Distributions

Evolutionary probability distributions were estimated using the antibody language model AntiBERTy [26] and the protein language model ESM2-650M [28]. When using AntiBERTy, we scored each variant by its pseudo-perplexity [30, 31]. When using ESM2-650M, we provided the wild-type sequence and computed, at each mutated position, the log-likelihood ratio of the mutant amino acid relative to the wild-type amino acid (referred to as wild-type marginal scoring in [32]). The score of a variant was defined as the sum of these log-likelihood ratios over its mutated positions. We report the absolute Spearman correlation |*ρ*| between each score and measured fitness. The AntiBERTy score was positively correlated with measured fitness in all five antibody landscapes, consistent with these landscapes measuring the fitness of somatic mutations away from the germline [10, 27], whereas the ESM2-650M score was negatively correlated with fitness only in the *β*-lactamase and Hu-1 landscapes. Following a convention similar to that of [33], variants with scores at or above the median score across the fitness landscape were designated as evolutionarily plausible. We defined the position-specific probability *p*_*i*_ as the fraction of evolutionarily plausible sequences carrying the mutant residue at position *i*, and the resulting vector forms the evolutionary measure ***p***. Because the score depends only on the sequence of a variant, landscapes that comprise the same variants (CR9114-H1 and CR9114-H3; CR6261-H1 and CR6261-H9) share the same set of evolutionarily plausible variants and therefore the same evolutionary measure. In the COV107-23 dataset, position two, as denoted in the original paper, is the F27X mutation, where X denotes a mutation to either V, L, or I. These three substitutions are treated as a single mutant state by summing their probabilities, and the log-likelihood ratio is computed relative to the wild-type residue. In the *β*-lactamase dataset, the first position analyzed is the nucleotide mutation g4205a (guanine to adenine at position 4205), so we computed the log-likelihood ratio over only the protein mutations and manually set *p*_1_ = 0.5. The resulting ***p*** distributions are detailed in the Supplementary Information.

To visualize the evolutionary manifold occupied by each protein (Fig. 1b), we computed mean-pooled language model embeddings from the final layer for each variant in each landscape. We then projected the embeddings using UMAP [29] with the cosine similarity metric. We randomly selected 500 evolutionarily plausible sequences to plot. To create Fig. 1d, we used the ***p*** distribution from Fig. 1c and visualized the Gram matrix up to second-order WHT basis functions evaluated on the evolutionary measure.

### Evaluation of Sparsity and Sparse Recovery

We evaluated eWHT and WHT on two sets of sequences. The first set comprised the evolutionarily plausible sequences defined above. The second set comprised experimentally high-fitness sequences, defined as those with measured fitness at or above the median measured fitness of the landscape, following a similar convention to [33].

To quantify spectral compactness, spectral coefficients were ranked by their absolute magnitudes, and cumulative variance explained (*R*^2^) was computed as more coefficients were retained on the evolutionarily plausible sequences. On the experimentally high-fitness sequences, mean squared error (MSE) was computed instead. For each landscape, we compared the number of coefficients, and the number of third-order and higher coefficients, required by eWHT and WHT to reach *R*^2^ = 0.95 (or an MSE of 0.05 on the experimentally high-fitness sequences). We computed the percentage reduction for each landscape and averaged these reductions across landscapes. Sparse recovery experiments were performed using Least Absolute Shrinkage and Selection Operator (LASSO) regression in both the eWHT and WHT domains. We trained our models on randomly sampled experimental measurements drawn from the evolutionarily plausible sequences. For each training subset, we selected the LASSO regularization value over [1 × 10^−5^, 5 × 10^−5^, 1 × 10^−4^, …, 0.1, 0.5] using five-fold cross-validation on the training data. We then tested our models on a held-out evaluation set of evolutionarily plausible sequences not used for training. Sparse recovery results represent averages over 10 independent training draws at each training size and five random seeds for evaluation pool construction.

### Interaction Visualization

We visualize the interactions required to achieve an *R*^2^ of 0.95 from Fig. 2b,c for Fig. 5. In Fig. 5a, we show the coefficients of the retained pairwise interactions for eWHT (upper triangle) and WHT (lower triangle), and compute the Spearman correlation over the pairwise interactions retained by both transforms. In Fig. 5b, we compute the *R*^2^ contribution of each interaction order as the sum, over the retained interactions of that order, of the incremental *R*^2^, defined as the change in the *R*^2^ curve when an interaction is added along the ranked coefficients of each transform. In Fig. 5c, for each interaction retained by either transform, we compute the difference between its incremental *R*^2^ under eWHT and under WHT and show the five interactions with the largest positive and the five with the largest negative difference. In Fig. 5d, we map the amino acids participating in the top five eWHT interactions from Fig. 5c onto the structures with PDB IDs 7LKA [35] for COV107-23, 4FQI [36] for CR9114-H1, and 3GBN [37] for CR6261-H1, coloring each amino acid by the highest interaction order in which it participates. As a co-crystal structure of CR9114 with H1 is not available, we use 4FQI, which is a structure of CR9114 bound to H5 hemagglutinin, following the original study [27]. For COV107-23, we denote the position number (e.g., in Fig. 5a, position 26 in G26E) as the position number of the amino acid in the canonical wild-type sequence. For CR9114-H1 and CR6261-H1, we follow the IMGT numbering used in the original paper [27].

### Synthetic Controls for the Genotype–Fitness Map and the Evolutionary Measure

To test whether the sparsity gain depends on the genotype–fitness map and on the evolutionary measure, we constructed five controls from each experimental landscape and repeated the compactness analysis on the evolutionarily plausible variants. Three controls altered the genotype–fitness map while holding the evolutionary measure and the evaluation variants fixed. The first control (shuffled fitness) permuted fitness values across genotypes. The second (global shuffle) permuted WHT coefficients across interaction orders, assigned each coefficient a random sign, and applied the inverse transform, preserving the distribution of coefficient magnitudes while reassigning them to different sets of sites. The third (banded shuffle) restricted the permutations to coefficients of the same order and likewise assigned random signs, also preserving the distribution of magnitudes within each order. Similarly, two controls altered the evolutionary measure while holding the genotype–fitness map and the evaluation variants fixed. The first (random measure) replaced the mutant frequency at each site with a value drawn uniformly at random from 0.05 to 0.95, and the second (measure shuffle) randomly permuted the mutant frequencies across sites. We generated each control using three random seeds (see Supplementary Information for the full results).

## Data availability

All fitness landscapes and ***p*** distributions used in this work are available at https://figshare.com/articles/dataset/eWHT_Data/32869433.

## Code availability

The code to reproduce all of our experiments is on GitHub at https://github.com/amirgroup-codes/eWHT_expts/. We additionally make our eWHT pip package software available at https://github.com/amirgroup-codes/ewht.

## Acknowledgements

This research was supported by the National Science Foundation (NSF) Graduate Research Fellowship Program (GRFP), the Exponential Electronics seed grant of the Institute for Matter and Systems (IMS) at Georgia Tech, Microsoft via GT Cloud Hub, Georgia Tech President’s Undergraduate Research Award, Georgia Tech Undergraduate Research Opportunities Program, and Georgia Institute of Technology start-up funds.

## Author contributions

D.T., K.T., and A.A. conceived the study. D.T. and K.T. developed the methodology, implemented the algorithms, and performed the experiments. D.T., K.T., and A.A. analyzed the results and wrote the manuscript. A.A. supervised the research.

## Competing interests

The authors declare no competing interests.

## Supplementary Information

### *p*-biased Analysis Details

The evolutionary Walsh–Hadamard transform (eWHT) used in the main text is the *p*-biased Walsh–Hadamard transform (*p*-WHT) [43], a classical orthonormal basis for functions defined on {−1, 1}^*L*^ under a product measure with the probability vector ***p*** = [*p*_1_, *p*_2_, …, *p*_*L*_], where *p*_*i*_ is the probability of the mutant state at site *i*. Because the results below hold for any ***p*** with 0 *< p*_*i*_ *<* 1, we use the terms *p*-WHT and eWHT interchangeably in this section. We write *π*_***p***_ for the product measure defined by ***p*** (Equation (1)) and ⟨·, ·⟩_***p***_ for the corresponding inner product (Equation (8)). Here, we briefly discuss relevant properties and derivations omitted from the main text.

### Properties of *p*-WHT Basis Functions

As mentioned in the main text, the *p*-WHT basis functions are of the form:

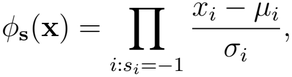

where *µ*_*i*_ = 1 − 2*p*_*i*_ and 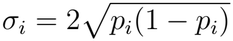. These terms are found as the mean and standard deviation of *x*_*i*_ under the evolutionary distribution of *x*_*i*_, which we write as:

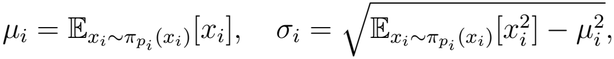

where 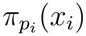 is the probability distribution of *x*_*i*_. By definition of the evolutionary probability measure, this is given as:

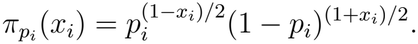

Therefore, plugging this into the above formulas of *µ*_*i*_ and *σ*_*i*_, *µ*_*i*_ can be found as:

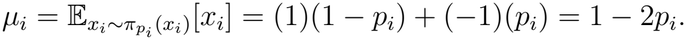

Similarly, *σ*_*i*_ can be found as:

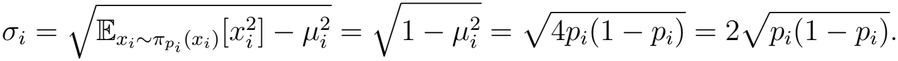

Next, we will prove an important theorem about the orthogonality of the basis functions. As an important preliminary fact, as seen in Section 1.2 of [43], *ϕ*_**1**_ = 1, where **1** is the length-*L* vector of ones.

#### Theorem 1.

*For all* ***p*** = [*p*_1_, *p*_2_, · · ·, *p*_*L*_] *with each p*_*i*_ ∈ (0, 1), *we have:*

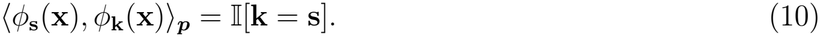

*Proof*. Let us start with the case where **s** = **k**. In this case, we have

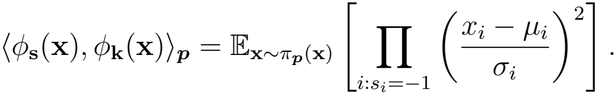

Since each *x*_*i*_ is independent, we can rewrite the expectation as:

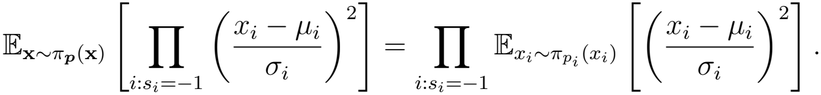

Seeing that 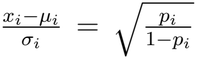 when *x*_*i*_ = 1 and 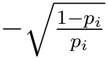 when *x*_*i*_ = −1, we can calculate the inner expectation as:

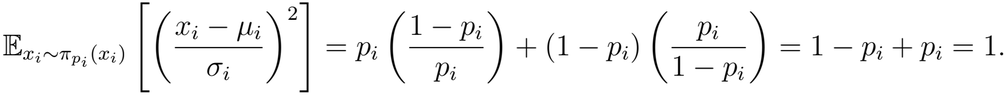

This gives us:

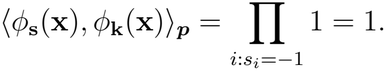

Next, let us consider the case where **s** ≠ **k**. In this case, we have:

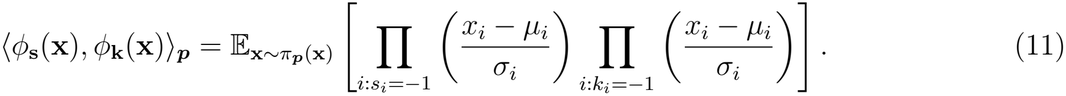

Since **s** ≠ **k**, we know there exists at least one index *j* such that *s*_*j*_ ≠ *k*_*j*_. Let the set *T* denote all indices that are −1 in **s** but not **k**, or −1 in **k** but not **s**. Then, we can rewrite Equation (11) as:

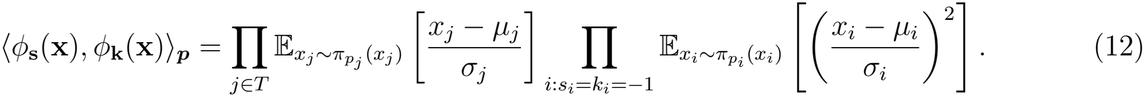

We can compute 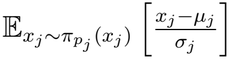 as:

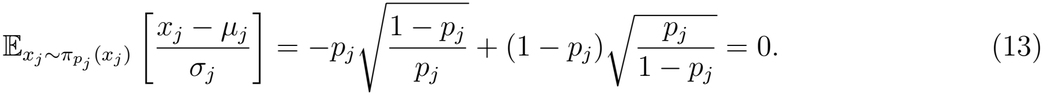

Since *T* is nonempty, the entire product in Equation (12) becomes 0, completing the proof.

Since the set of basis functions spans the set of all fitness functions *f* : {−1, 1}^*L*^ → R, every such function has a unique *p*-WHT decomposition [43].

### Matrix Multiplication Formulation of the *p*-WHT

Note that throughout this section, following classical Walsh–Hadamard notation, we define index 1 associated with *p*_1_ as the least significant bit and index *L* associated with *p*_*L*_ as the most significant bit. In practice, our implementation uses the opposite convention for convenience, but the theoretical results are unchanged by this. Given a length-2^*L*^ vector of fitness values **f**, the *p*-WHT coefficients 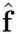 can be computed as:

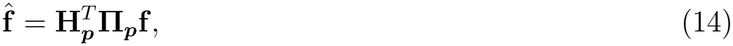

where 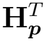 is the transpose of **H**_***p***_ and **Π**_***p***_ = diag (*π*_***p***_(**x**)) is the diagonal matrix of the probabilities of all sequences under *π*_***p***_. Both 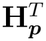 and **Π**_***p***_ can be decomposed using the Kronecker product: 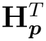 as 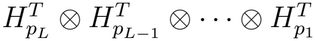, where

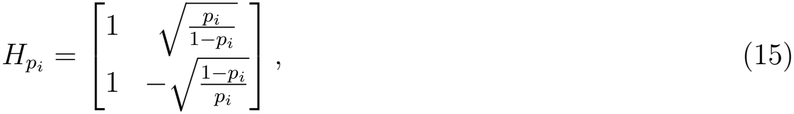

and **Π**_***p***_ as 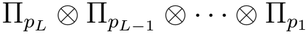, where

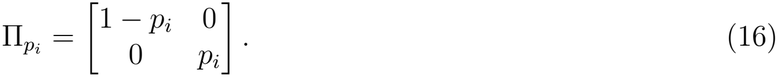

Clearly, the diagonal matrix **Π**_***p***_ represents the expectation weighting of the *p*-biased inner product, while the columns of **H**_***p***_ represent the basis functions *ϕ*_**s**_(**x**) evaluated over all 2^*L*^ values for **s** and **x**. If all *p*_*i*_ are set to 0.5, this formulation reduces to that of the WHT, and **H**_***p***_ becomes the Hadamard matrix.

Next, we provide a short corollary to show that in matrix form, the *p*-WHT is invertible. Note that **Π**_***p***_ is only applied in the forward transform, since the reverse transform does not have an inner product.

#### Corollary 1.

*For all* ***p*** = [*p*_1_, *p*_2_, · · ·, *p*_*L*_] *with each p*_*i*_ ∈ (0, 1) *for all i, we have*

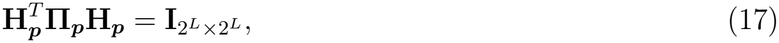

*where* 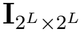 *is the* 2^*L*^ × 2^*L*^ *identity matrix*.

*Proof*. Since each row of 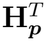 is a basis function *ϕ*_**s**_ evaluated at all 2^*L*^ values of **x**, and each column of **H**_***p***_ is the same, we can see that:

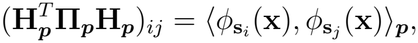

where **s**_*i*_ and **s**_*j*_ are the vectors in {−1, 1}^*L*^ corresponding to the basis functions at indices *i* and *j*. By Theorem 1, we know the right-hand side will only equal 1 when *i* = *j*, and otherwise equals 0. This means that:

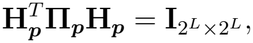

concluding the proof.

### FFT Implementation of the *p*-WHT

Since the *p*-WHT has a structure similar to the WHT, it can be computed using an algorithm that runs in O(*N* log *N*), where *N* = 2^*L*^. The structure of this algorithm mirrors that of the fast Walsh–Hadamard transform and reduces to it if all *p*_*i*_ = 0.5. The forward fast *p*-WHT (abbreviated as *p*FWHT) is detailed in Algorithm 1, and the inverse (abbreviated as I*p*FWHT) is detailed in Algorithm 2. It can be seen that these algorithms are both *O*(*N* log *N*) since there are *L* = log_2_(2^*L*^) outer loops over *i*, and within each outer loop we have 2^*L*^*/*2^*i*^ inner loops over *j*, within which we have 2^*i*−1^ inner loops over *k*. This gives a complexity of:

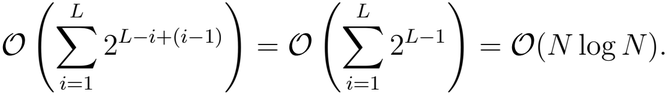

Next, we will prove that Algorithm 1 matches the matrix multiplication of the *p*-WHT.

#### Theorem 2.

*For all* ***p*** = [*p*_1_, *p*_2_, · · ·, *p*_*L*_] *with each p*_*i*_ ∈ (0, 1), *we have*

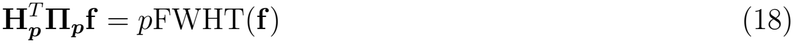

*for all* 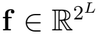.

*Proof*. We proceed by induction on *L*, with a base case of *L* = 1. In this case, each loop (*i, j, k*) of Algorithm 1 runs once, giving an output of

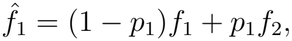

and

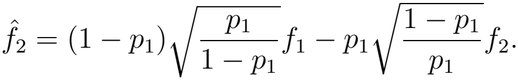

Clearly, this matches the result of multiplying **f** by 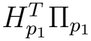. Now, let us assume that for some arbitrary *k* ∈ N with arbitrary ***p*** = [*p*_1_, *p*_2_, · · ·, *p*_*k*_] and 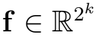, it holds that

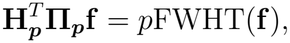

where

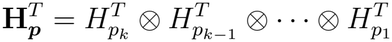

and

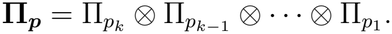

Now, define an arbitrary 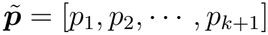, and an arbitrary 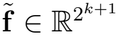. To prove the theorem, we will prove the inductive step, which is that

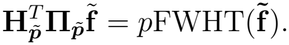

If we run Algorithm 1 on 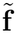 until *i* = *k*, then the resulting output will have applied the *p*FWHT on ***p*** individually to the top and bottom half of 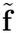. Splitting 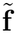 into **a** and **b**, we can write this as 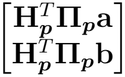, where we use the induction hypothesis to equate the *p*FWHT to the matrix multiplication. Running the last step of the algorithm with *i* = *k* + 1 on the given vector, we see that the output is given as:

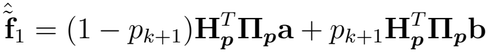

and

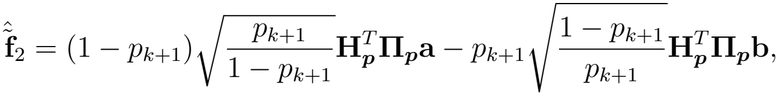

where 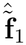 and 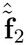 represent the first and second halves of the output 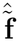. This can be rewritten as a matrix equation:

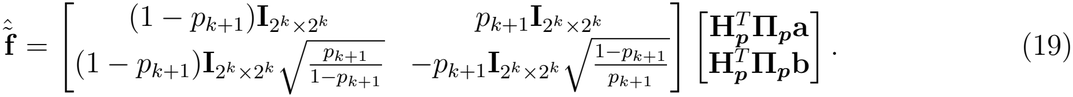

This can again be rewritten using the Kronecker product as:

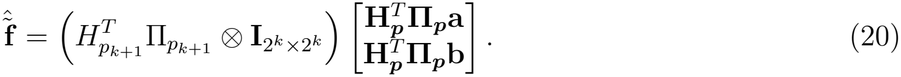

This can be simplified further as:

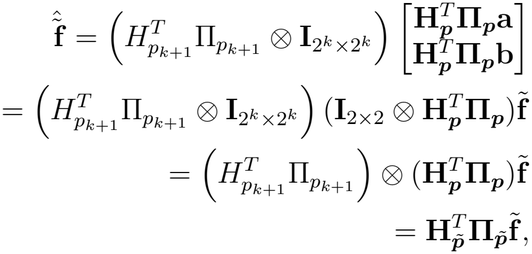

where the third line comes from the mixed-product property of Kronecker products. This proves the inductive step and the theorem.

As a simple consequence of this theorem, we can prove that Algorithm 2 matches the matrix multiplication of the inverse *p*-WHT with the same proof strategy.

#### Corollary 2.

*For all* ***p*** = [*p*_1_, *p*_2_, · · ·, *p*_*L*_] *with each p*_*i*_ ∈ (0, 1), *we have*

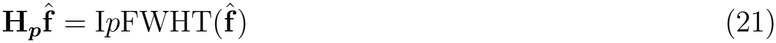

*for all* 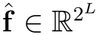.

*Proof*. We similarly proceed by induction, again starting with a base case of *L* = 1. This would give us:

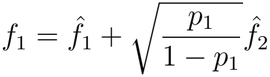

and

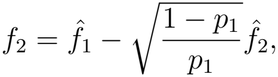

clearly matching the result of multiplying 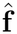 by 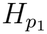. For the induction hypothesis, we again assume that for some *k* ∈ N and ***p*** = [*p*_1_, *p*_2_, · · ·, *p*_*k*_], that:

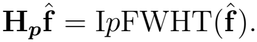

Now, define an arbitrary 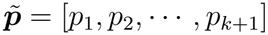, and an arbitrary 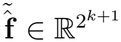. Running Algorithm 2 on 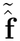 until *i* = *k* will apply the I*p*FWHT to the top and bottom half of 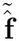. If we again split 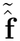 into a top and bottom **a** and **b**, we can write this as 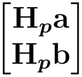, again using the induction hypothesis. Running the last step of the algorithm gives us:

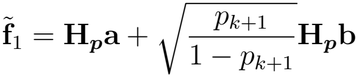

and

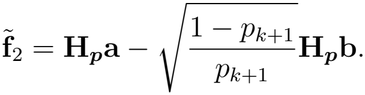

We can write this in matrix multiplication form as:

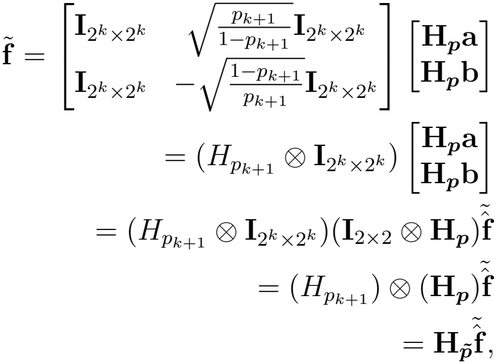

completing the proof.

### Converting Coefficients from a Uniform to a Non-uniform Measure

In this section, we discuss why a non-uniform measure converts uniform basis coefficients into a sum of non-uniform basis coefficients of equal or lower interaction order, as seen in Equation (18) of [25]. Let us write the standard Walsh–Hadamard basis function as:

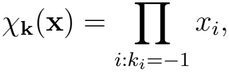

which is the special case of the *p*-WHT basis function when *p*_*i*_ = 0.5 for all *i*, so that *µ*_*i*_ = 0 and *σ*_*i*_ = 1. Next, we decompose *χ*_**k**_ in the *p*-WHT basis:

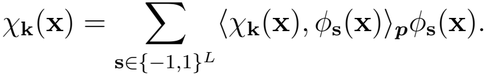

We can find the inner product ⟨*χ*_**k**_(**x**), *ϕ*_**s**_(**x**)⟩_***p***_ as:

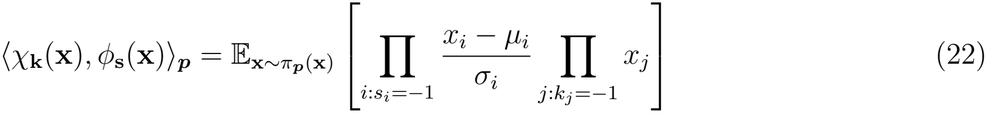

Additionally, let us define the sets:

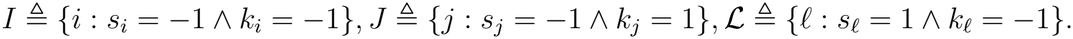

We can then rewrite Equation (22) as:

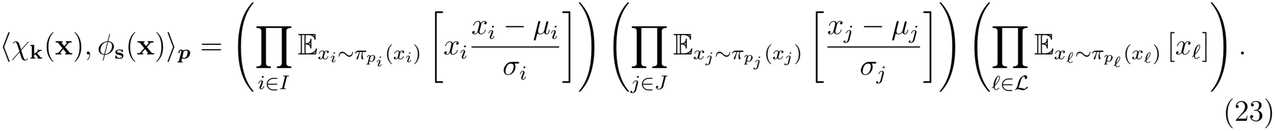

As seen in Equation (13), the product over *J* becomes zero, meaning that ⟨*χ*_**k**_(**x**), *ϕ*_**s**_(**x**)⟩_***p***_ = 0 unless *J* = ∅. This implies that a Walsh–Hadamard coefficient **k** can be represented in terms of *p*-WHT basis functions indexed by subsets of **k**.

### Sparse Recovery Details

Our sparse recovery results are based on compressed sensing [44], an area of signal processing that uses sparse representations of signals in othogonal bases to solve underdetermined linear systems of the form:

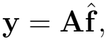

where, in this case, **y** ∈ ℝ^*m*^ represents the fitness measurements, 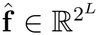 represents the sparse vector of *p*-WHT coefficients, and 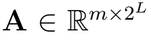 represents the matrix of basis values *ϕ* that map from the *p*-WHT representation to the fitness representation. We make the assumption that 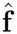 is *sparse*, meaning only a small subset of its entries are non-zero, allowing us to solve a compressed sensing problem of this form. The rows of **A** are found as the rows of **H**_***p***_ such that 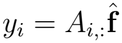, where *A*_*i*,:_ is the *i*^th^ row of **A**.

## Supplementary Figures

### Algorithm 1

Fast *p*-WHT (*p*FWHT)

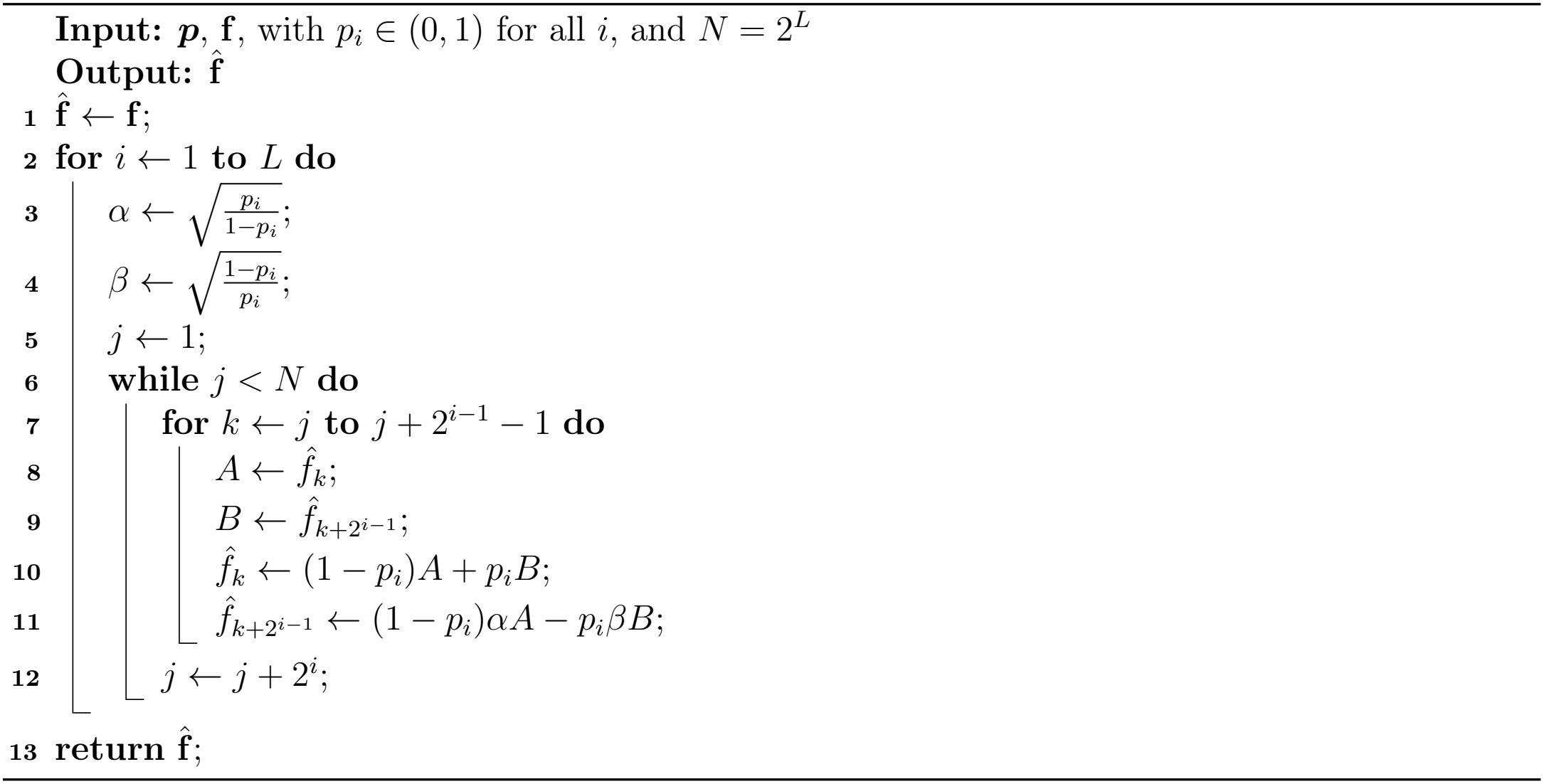

### Algorithm 2

Inverse Fast *p*-WHT (I*p*FWHT)

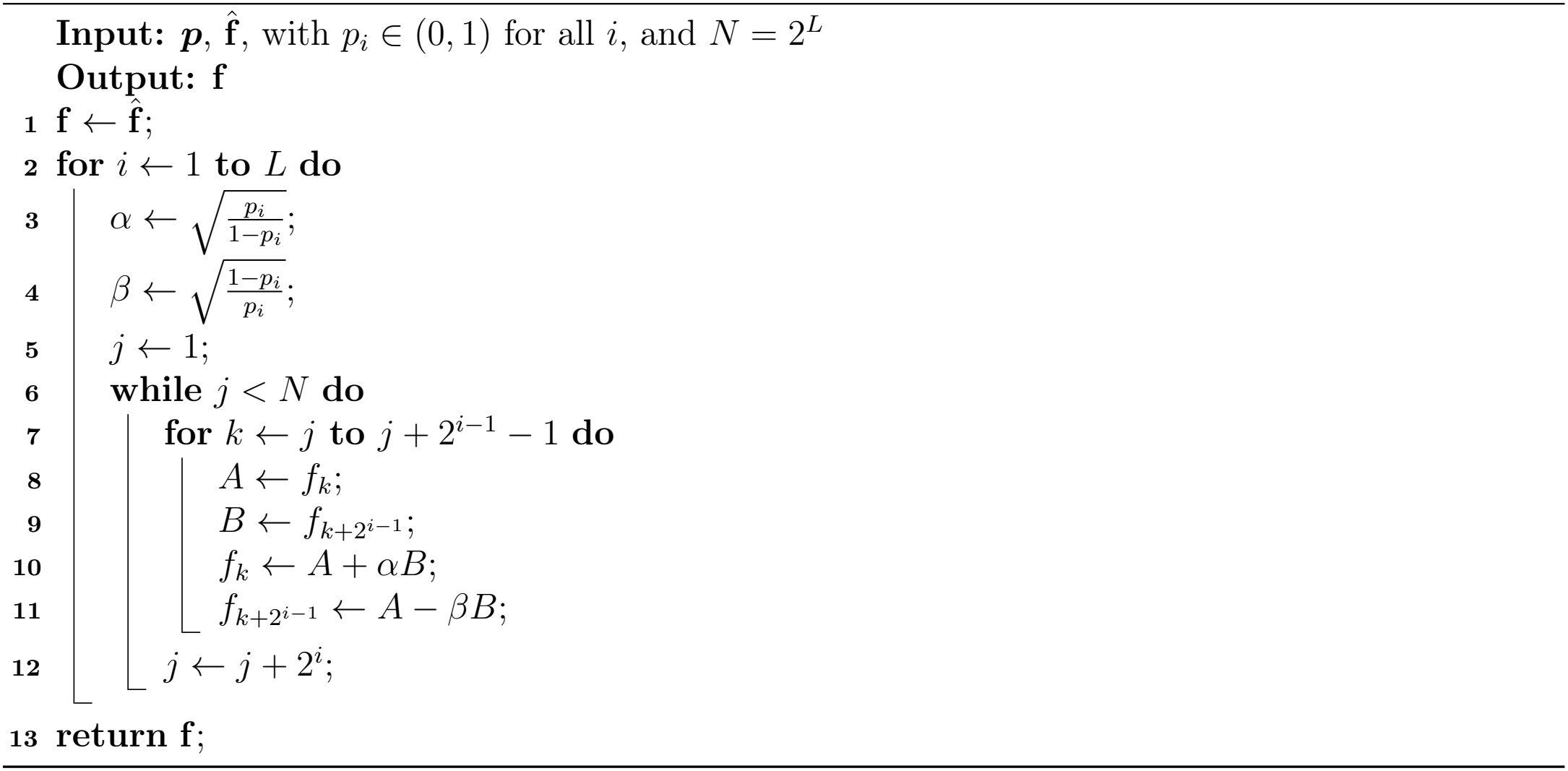

**Figure S1:**
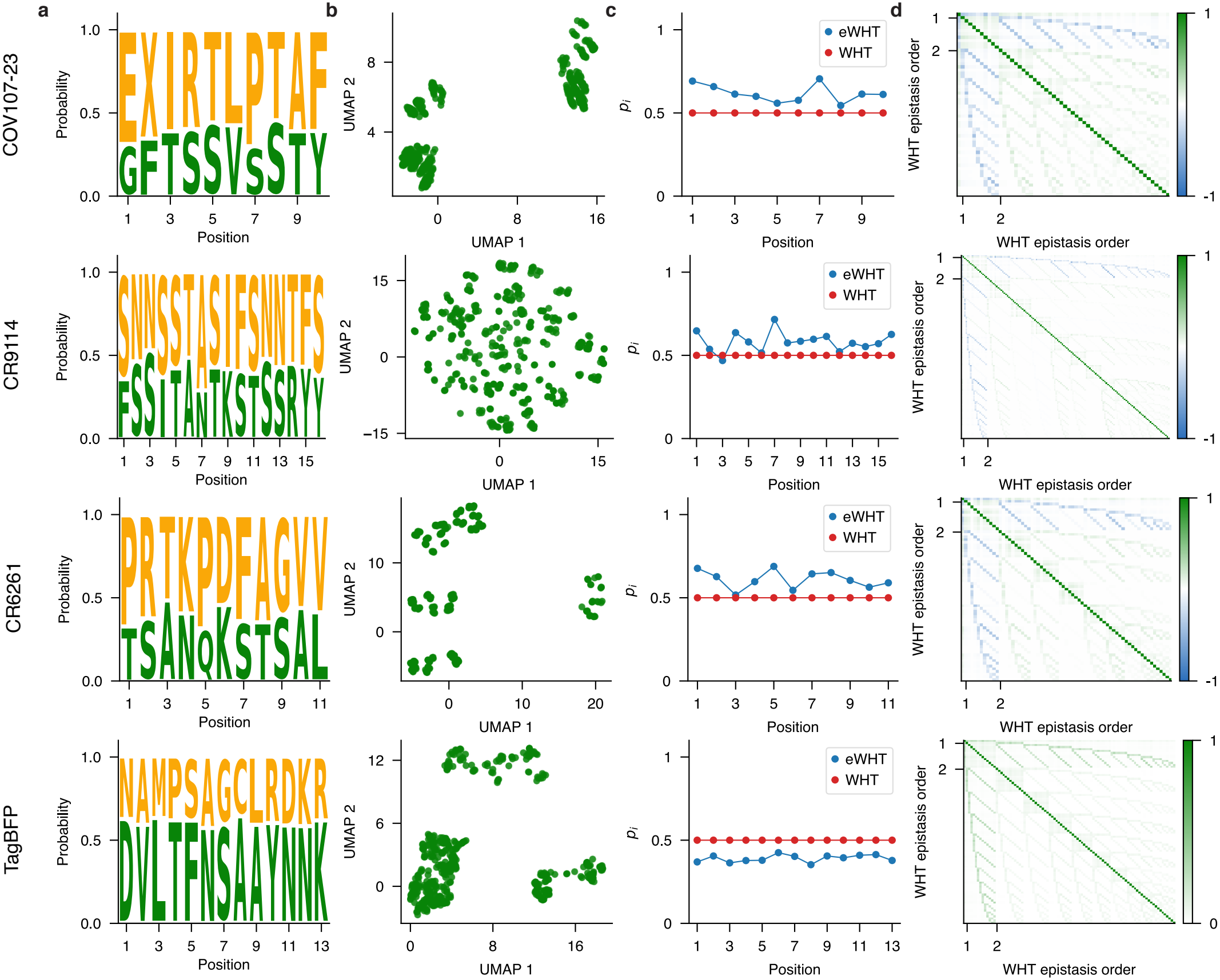
Evolutionary distributions of the (top to bottom) COV107-23, CR9114 (H1 and H3), CR6261 (H1 and H9), and TagBFP landscapes. Landscapes that comprise the same variants share the same evolutionary measure. **a** Wild-type (green) and mutant (orange) amino acid frequencies at each site among the evolutionarily plausible sequences. **b** UMAP projections of language model embeddings of the evolutionarily plausible sequences (AntiBERTy for the antibody landscapes and ESM2-650M for TagBFP). **c** Mutant frequency *p*_*i*_ at each site *i*, defined as the fraction of evolutionarily plausible sequences carrying the mutant residue. The classical Walsh–Hadamard transform (WHT) implicitly assumes *p*_*i*_ = 0.5 at every site, corresponding to a uniform distribution over sequence space. **d** Gram matrix of the classical WHT basis functions through second order, evaluated under the evolutionary measure defined by the frequencies in **c**. The non-zero off-diagonal entries show that the classical WHT basis loses its orthonormality under the evolutionary distribution.

**Figure S2:**
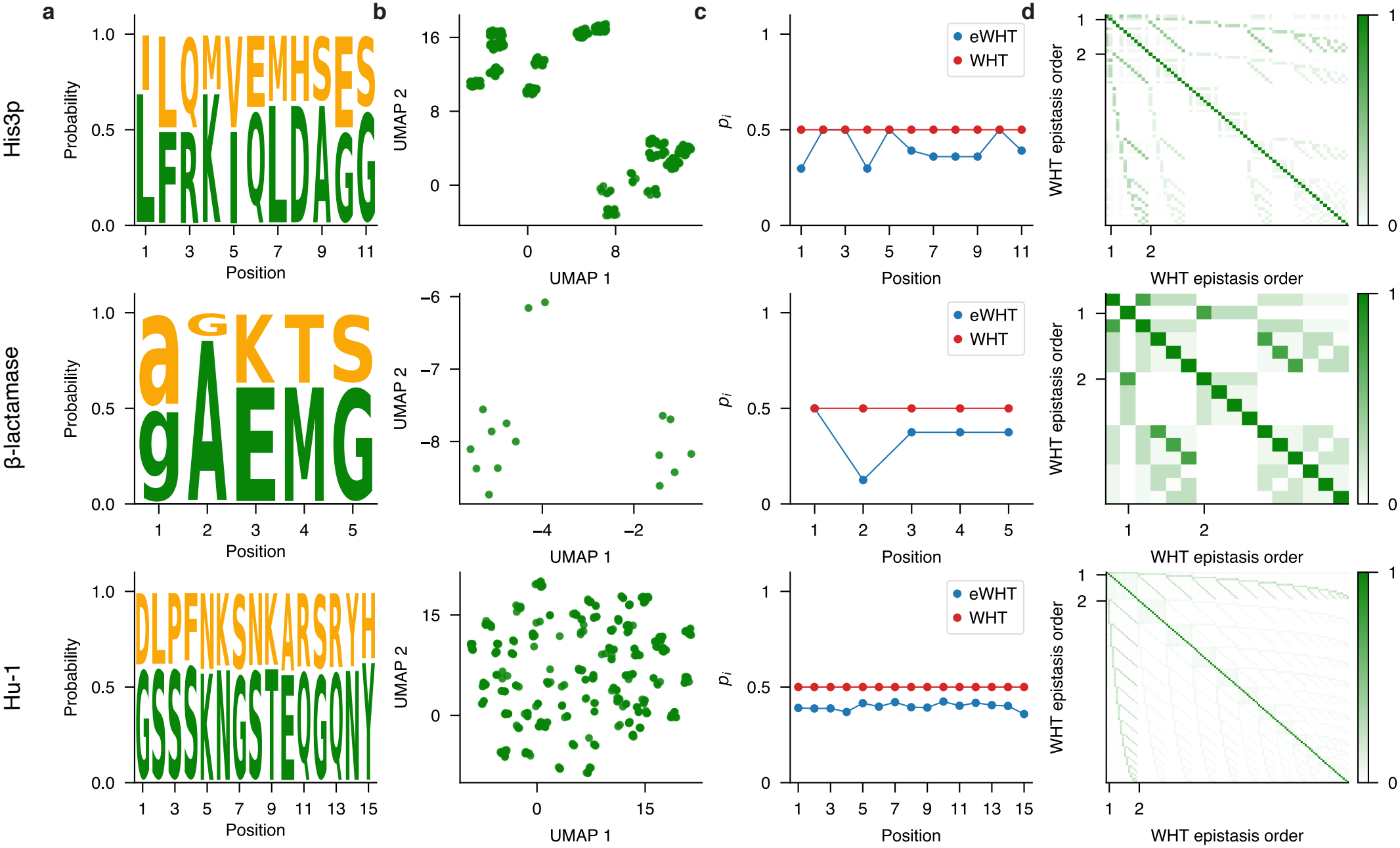
Evolutionary distributions of the (top to bottom) His3p, *β*-lactamase, and Hu-1 landscapes. Panels **a**–**d** are as in Fig. S1, with ESM2-650M embeddings in panel **b**. In the *β*-lactamase landscape, the first site is the nucleotide mutation from guanine (g) to adenine (a) at position 4205, for which *p*_1_ was set to 0.5.

**Figure S3:**
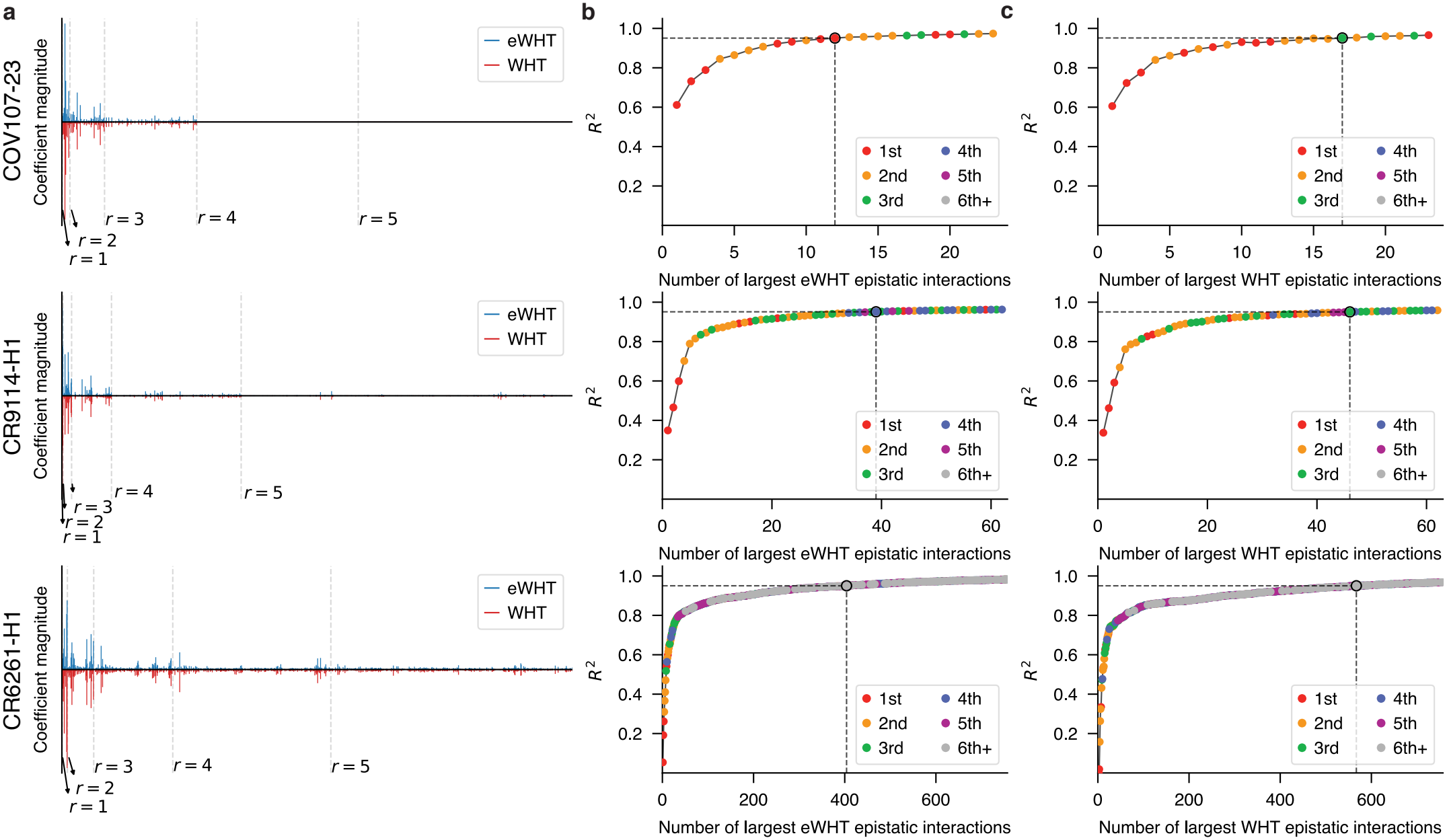
Comparison of fitness function spectra under eWHT and WHT of (top to bottom) COV107-23, CR9114-H1, and CR6261-H1 evaluated on evolutionarily plausible sequences. **a** Magnitude of spectral coefficients of eWHT (upper plot, in blue) compared to WHT (lower plot, in red). Dashed lines separate orders of epistatic interactions, showing up to 5^th^-order interactions. **b** *R*^2^ plot of evolutionarily plausible sequences against number of eWHT epistatic interactions, arranged by largest interactions. Colors show the order of epistasis. The dashed lines mark the smallest number of interactions for which *R*^2^ ≥ 0.95. **c** *R*^2^ plot of evolutionarily plausible sequences against number of WHT epistatic interactions.

**Figure S4:**
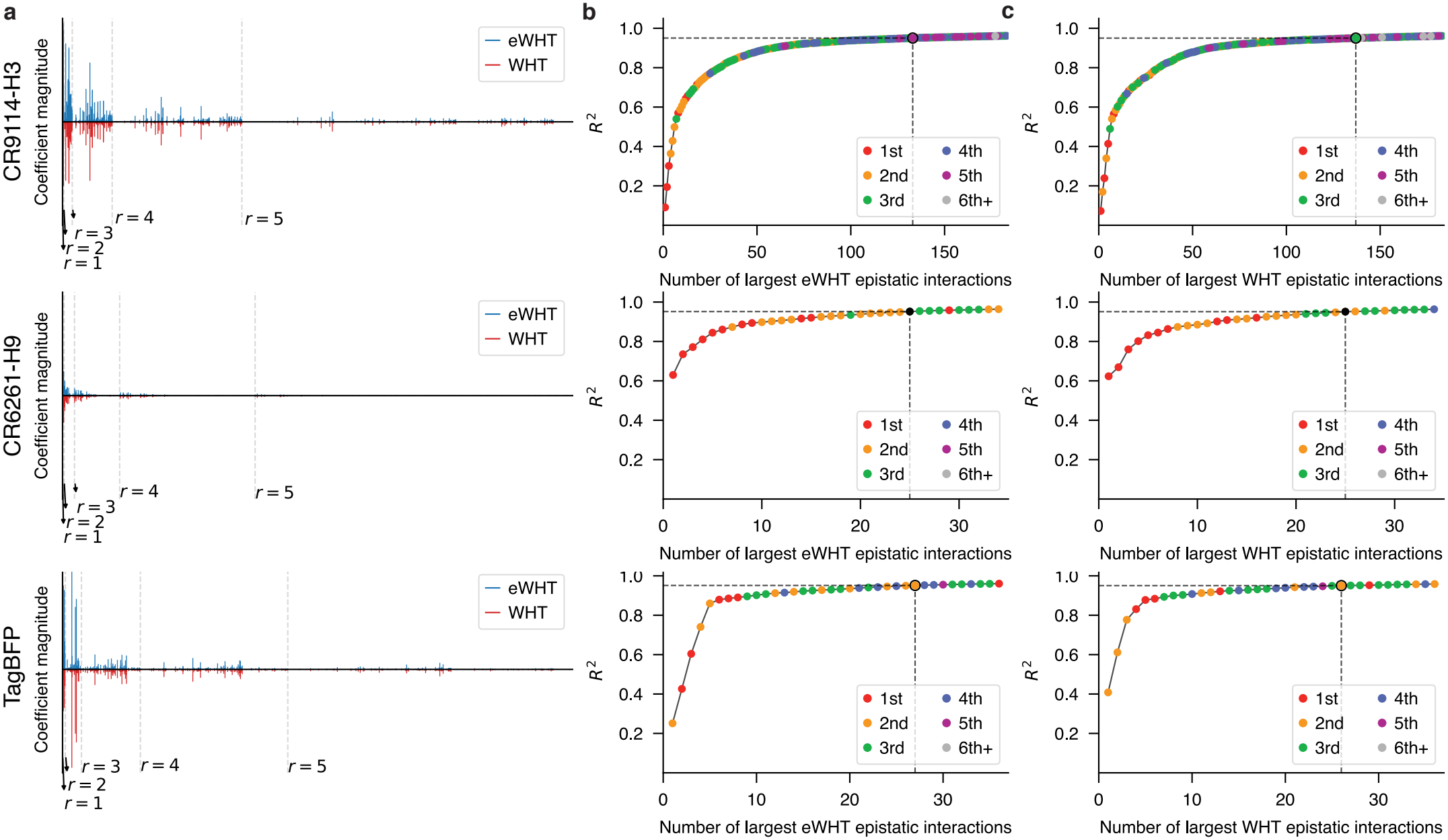
Comparison of fitness function spectra under eWHT and WHT of (top to bottom) CR9114-H3, CR6261-H9, and TagBFP evaluated on evolutionarily plausible sequences. Panels **a**–**c** are as in Fig. S3.

**Figure S5:**
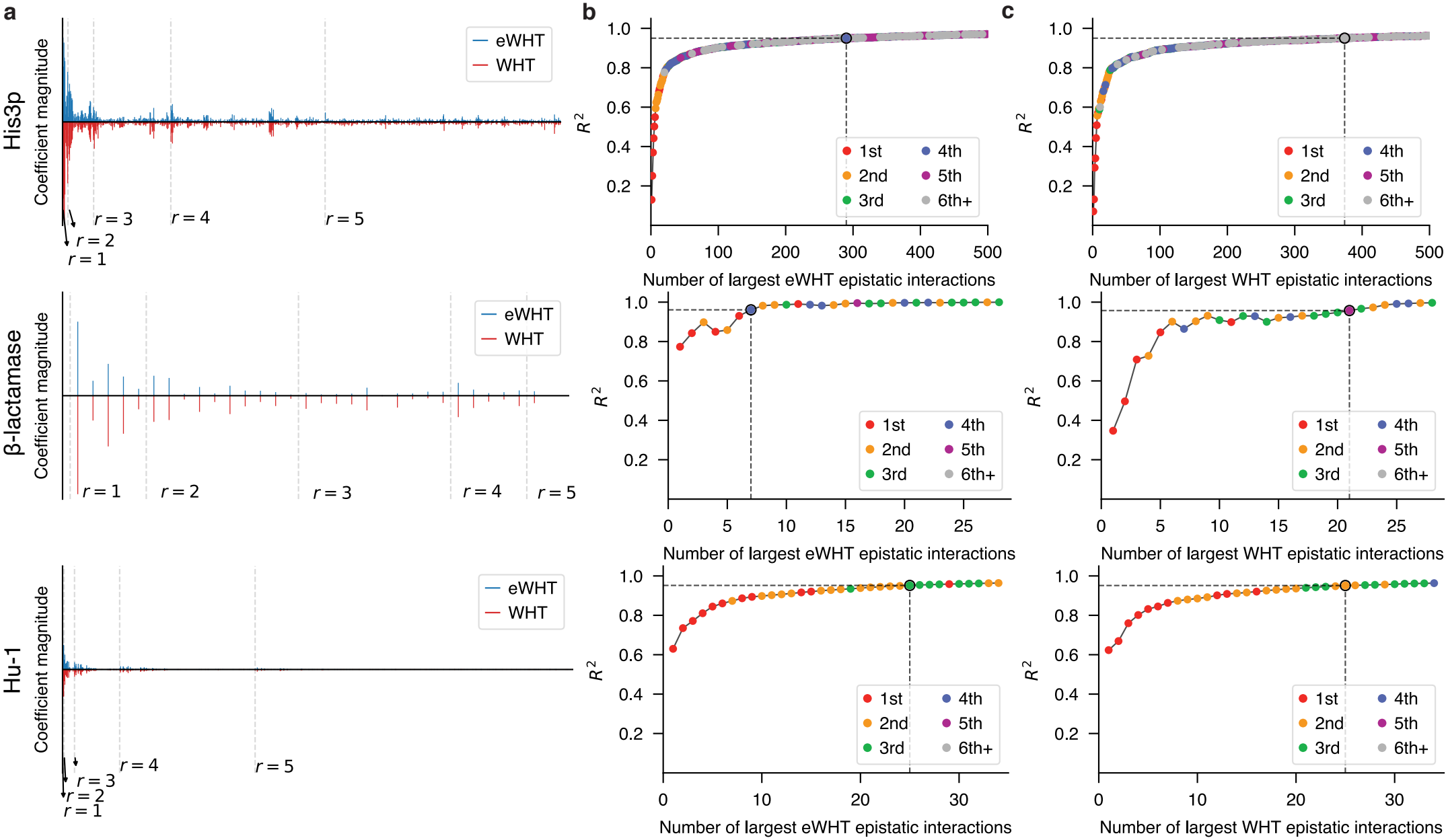
Comparison of fitness function spectra under eWHT and WHT of (top to bottom) His3p, *β*-lactamase, and Hu-1 evaluated on evolutionarily plausible sequences. Panels **a**–**c** are as in Fig. S3.

**Figure S6:**
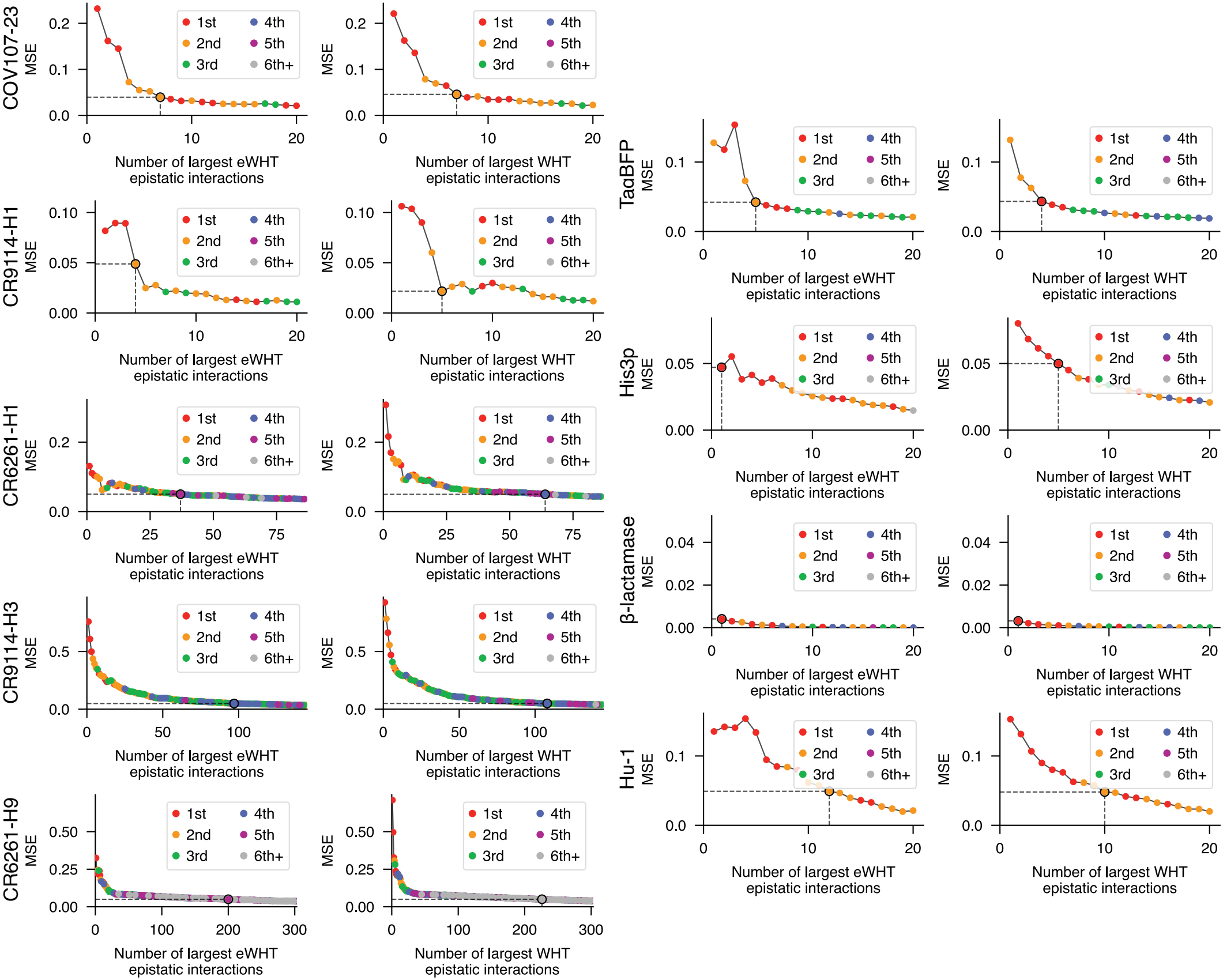
Comparison of fitness function spectra under eWHT and WHT of (top to bottom, left to right) COV107-23, CR9114-H1, CR6261-H1, CR9114-H3, CR6261-H9, TagBFP, His3p, *β*-lactamase, and Hu-1. Mean squared error (MSE) plot of experimental high-fitness sequences against number of eWHT and WHT epistatic interactions, arranged by largest interactions. Colors show the order of epistasis. The black dot and dashed lines mark the smallest number of interactions for which MSE ≤ 0.05.

**Figure S7:**
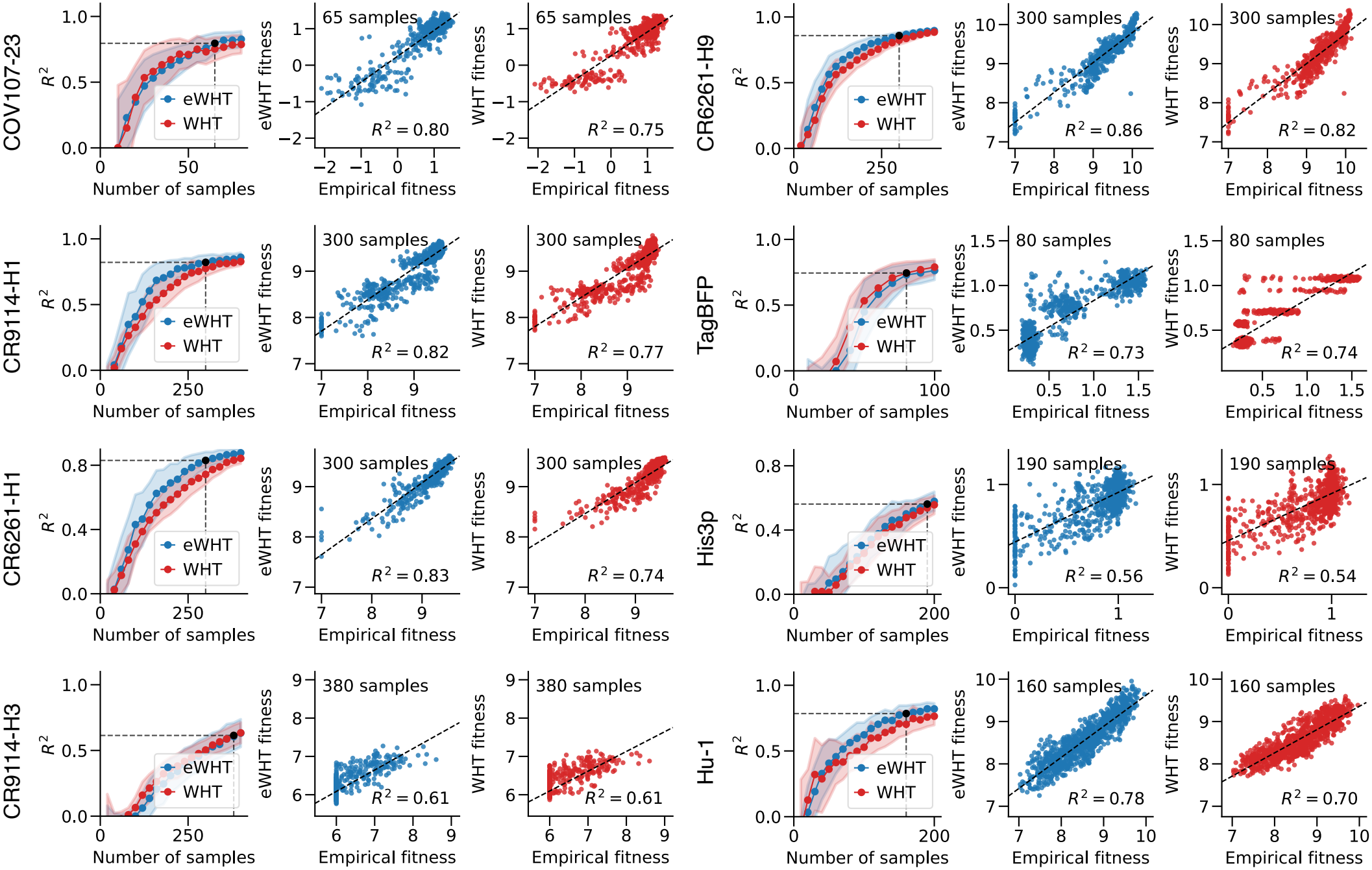
Estimation of fitness with compressed sensing with eWHT and WHT evaluated on held-out evolutionarily plausible sequences. The left-most plot for each protein displays the *R*^2^ of evolutionarily plausible sequences at a range of training set sizes. The middle plot displays the reconstruction of fitness from estimated eWHT epistatic interactions. The right-most plot displays reconstruction of fitness from estimated WHT epistatic interactions. Reconstruction plots were chosen based on 95% of the average *R*^2^ at the maximum number of samples tested. As mentioned in the main text, the *β*-lactamase protein is excluded due to containing only 32 variants.

**Figure S8:**
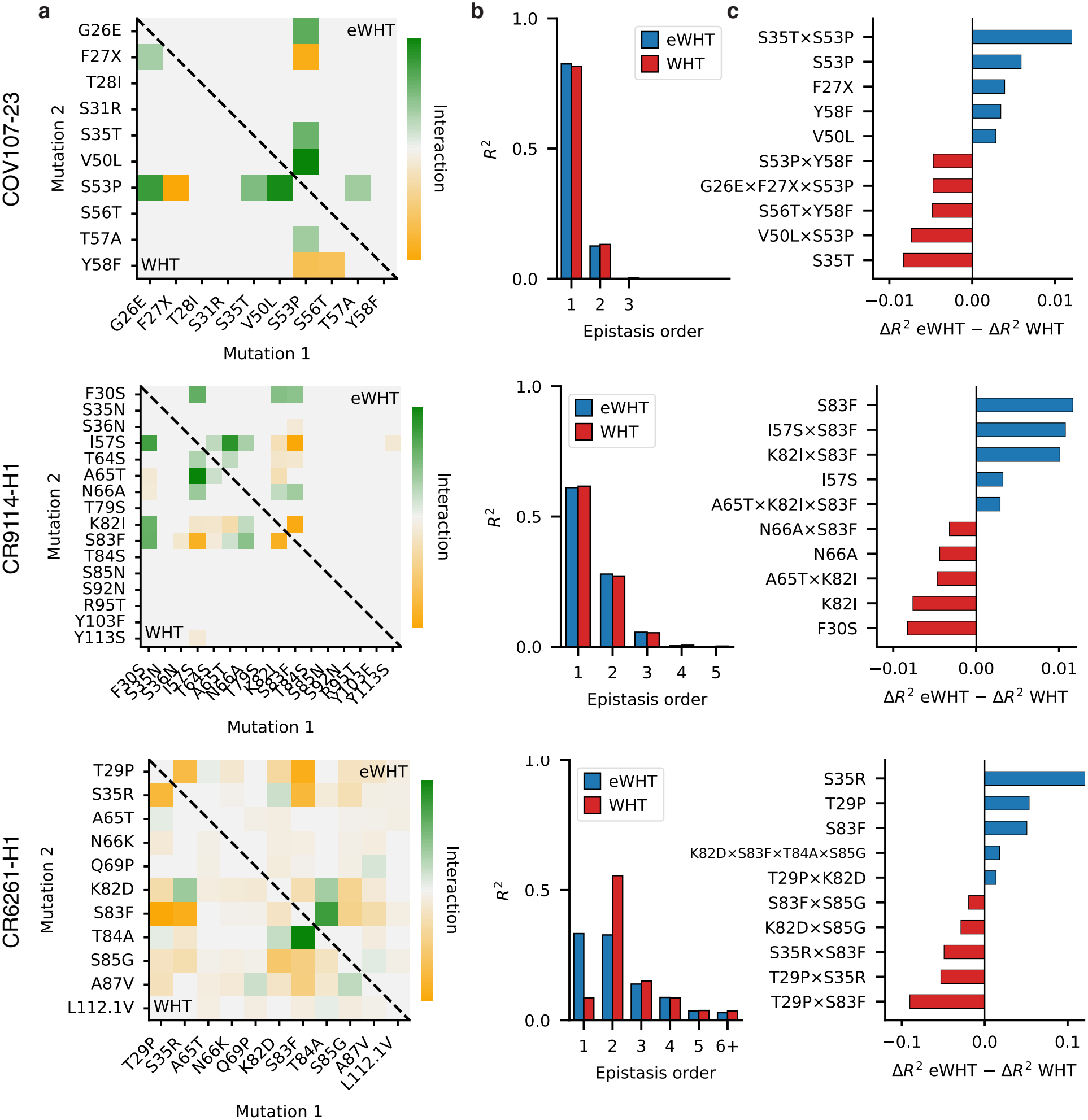
Terms retained by eWHT and WHT in the (top to bottom) COV107-23, CR9114-H1, and CR6261-H1 landscapes when each expansion is truncated at *R*^2^ = 0.95 among evolutionarily plausible variants (Fig. S3b,c). **a** Pairwise coefficients retained by eWHT (upper triangle) and WHT (lower triangle). The X mutation in COV107-23 denotes a mutation to V, L, or I. **b** *R*^2^ contributed by the retained terms, stratified by interaction order. **c** The five terms with the largest increase in incremental *R*^2^ under eWHT (positive) and under WHT (negative), shown as 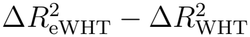. Incremental *R*^2^ is the change in the variance-explained curve of Fig. S3b,c when a term is added. Panels **a**–**c** correspond to panels **a**–**c** of Fig. 5, which additionally maps residues onto structures.

**Figure S9:**
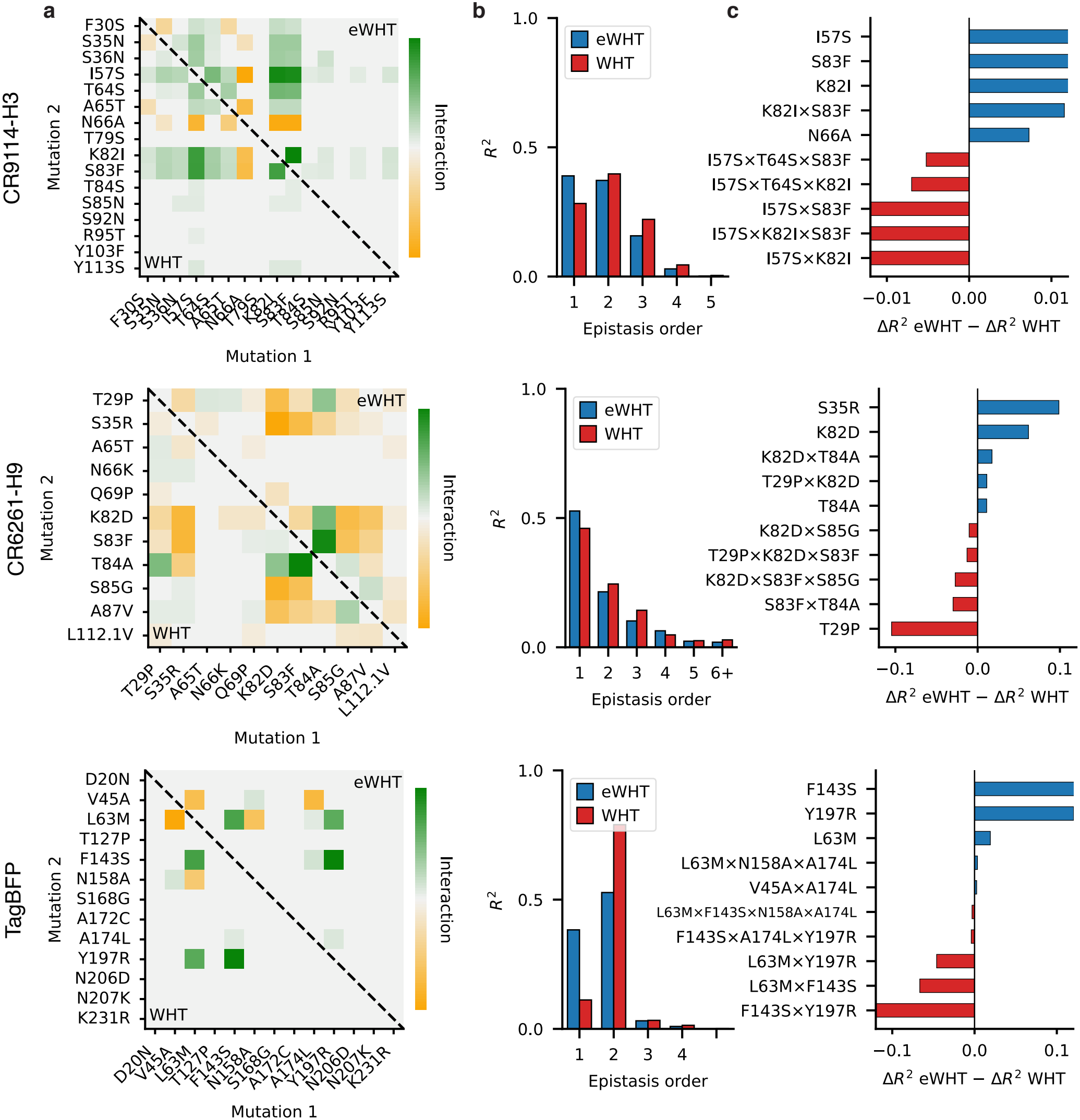
Terms retained by eWHT and WHT in the (top to bottom) CR9114-H3, CR6261-H9, and TagBFP landscapes when each expansion is truncated at *R*^2^ = 0.95 among evolutionarily plausible variants (Fig. S4b,c). Panels **a**–**c** are as in Fig. S8.

**Figure S10:**
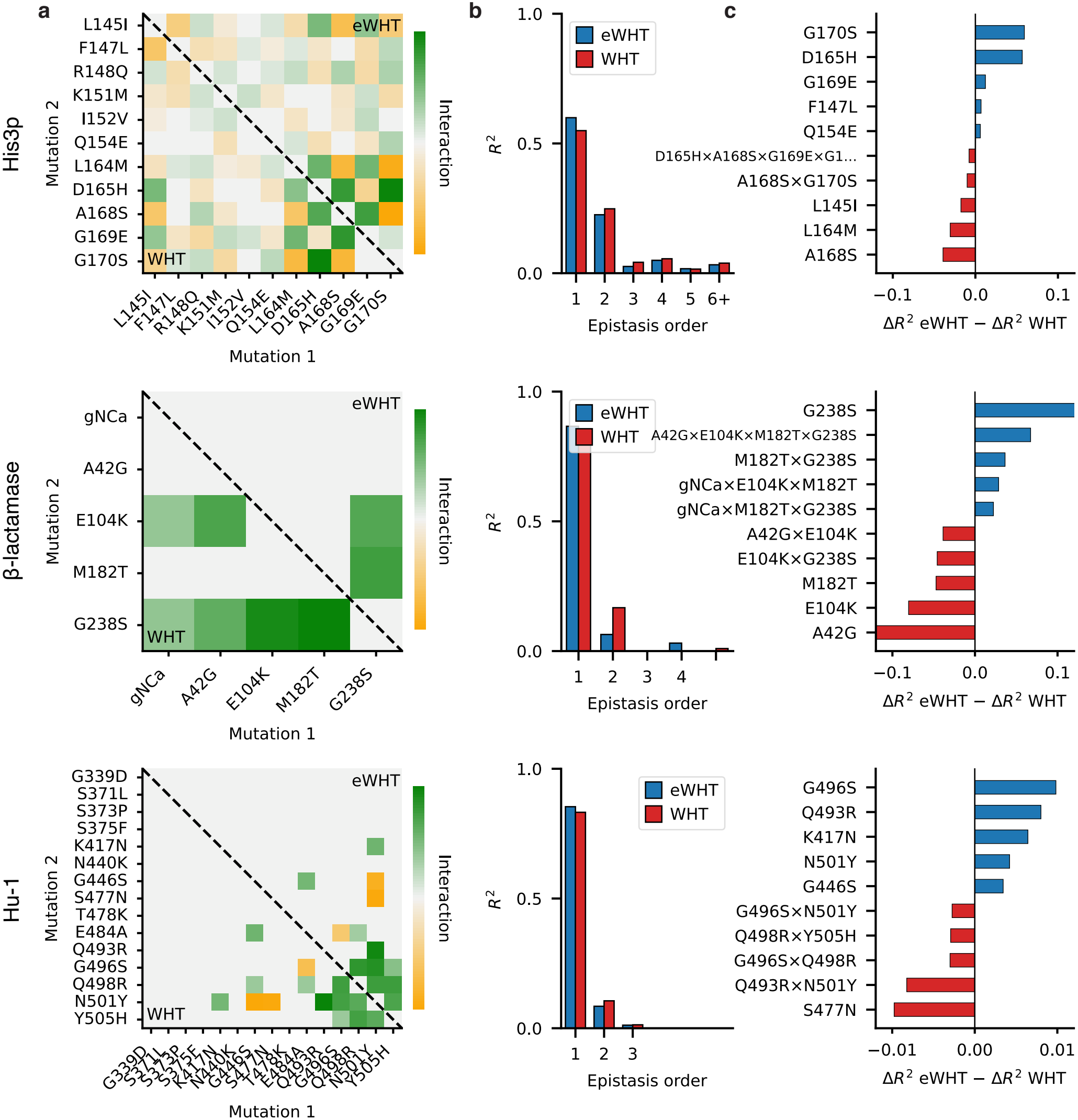
Terms retained by eWHT and WHT in the (top to bottom) His3p, *β*-lactamase, and Hu-1 landscapes when each expansion is truncated at *R*^2^ = 0.95 among evolutionarily plausible variants (Fig. S5b,c). Panels **a**–**c** are as in Fig. S8.

**Figure S11:**
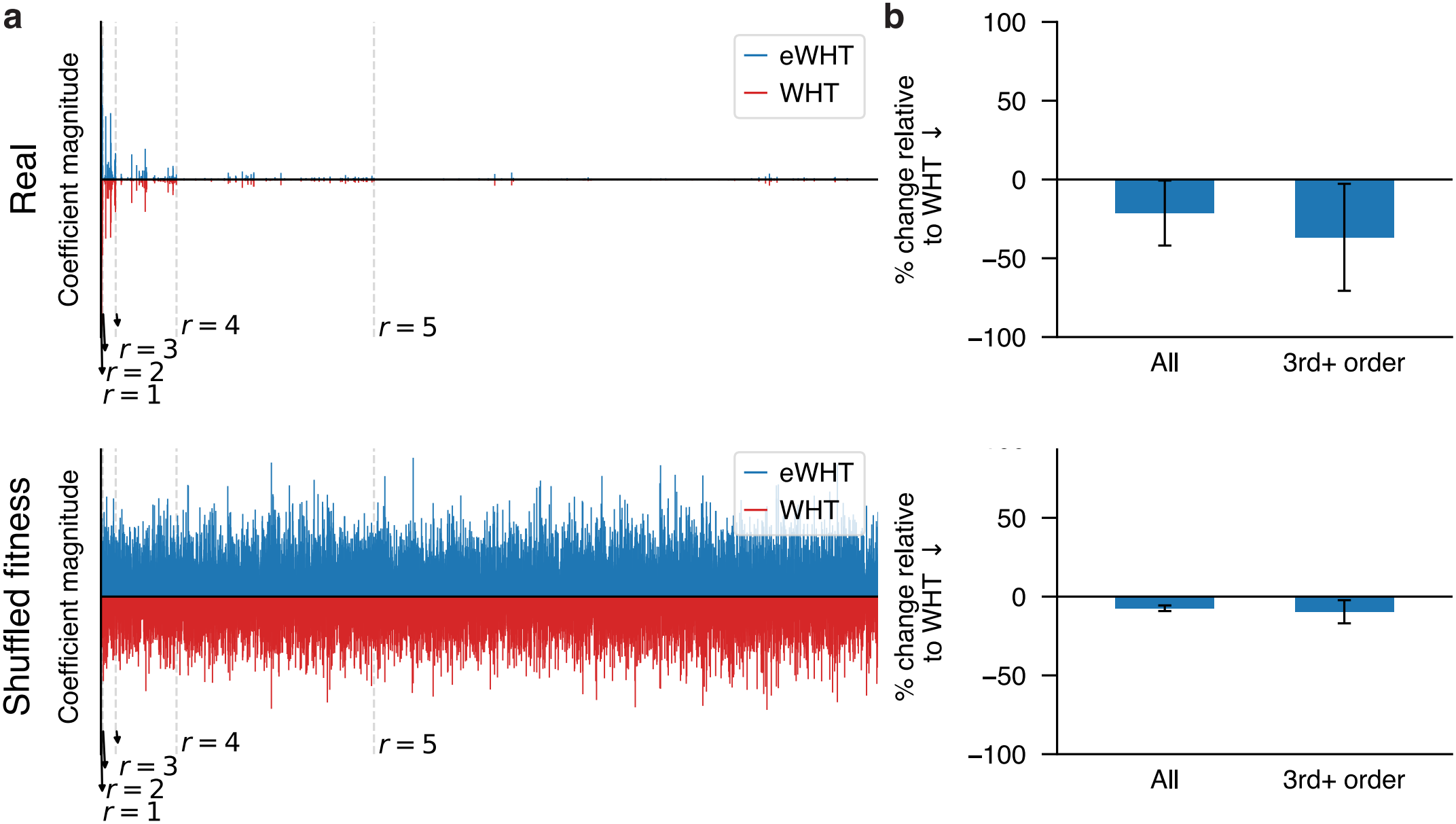
Permuting fitness values weakens the advantage of eWHT. Comparison of eWHT and WHT on the experimental fitness landscapes (real) and on the shuffled fitness synthetic landscape, in which the fitness values of all sequences are randomly permuted across genotypes (see Methods). In the shuffled fitness landscape, the evolutionary measure and evaluation sequences are those of the experimental landscape, and only the sequence-to-function mapping differs. **a** Spectral coefficients of eWHT (blue, upward) and WHT (red, downward) for the CR9114-H1 landscape and, for the shuffled fitness landscape, one random seed. **b** Percentage reduction in the number of coefficients (All) and in the number of third-order and higher coefficients (3rd+ order) required by eWHT relative to WHT to reach *R*^2^ = 0.95 on the evolutionarily plausible sequences, computed for each landscape and averaged across the nine landscapes (synthetic values are first averaged over three random seeds, and error bars show the standard deviation across landscapes). Blue bars indicate that eWHT required fewer coefficients than WHT, and red bars indicate that eWHT required more. The average reduction is larger on the experimental landscapes than on the shuffled fitness landscapes.

**Figure S12:**
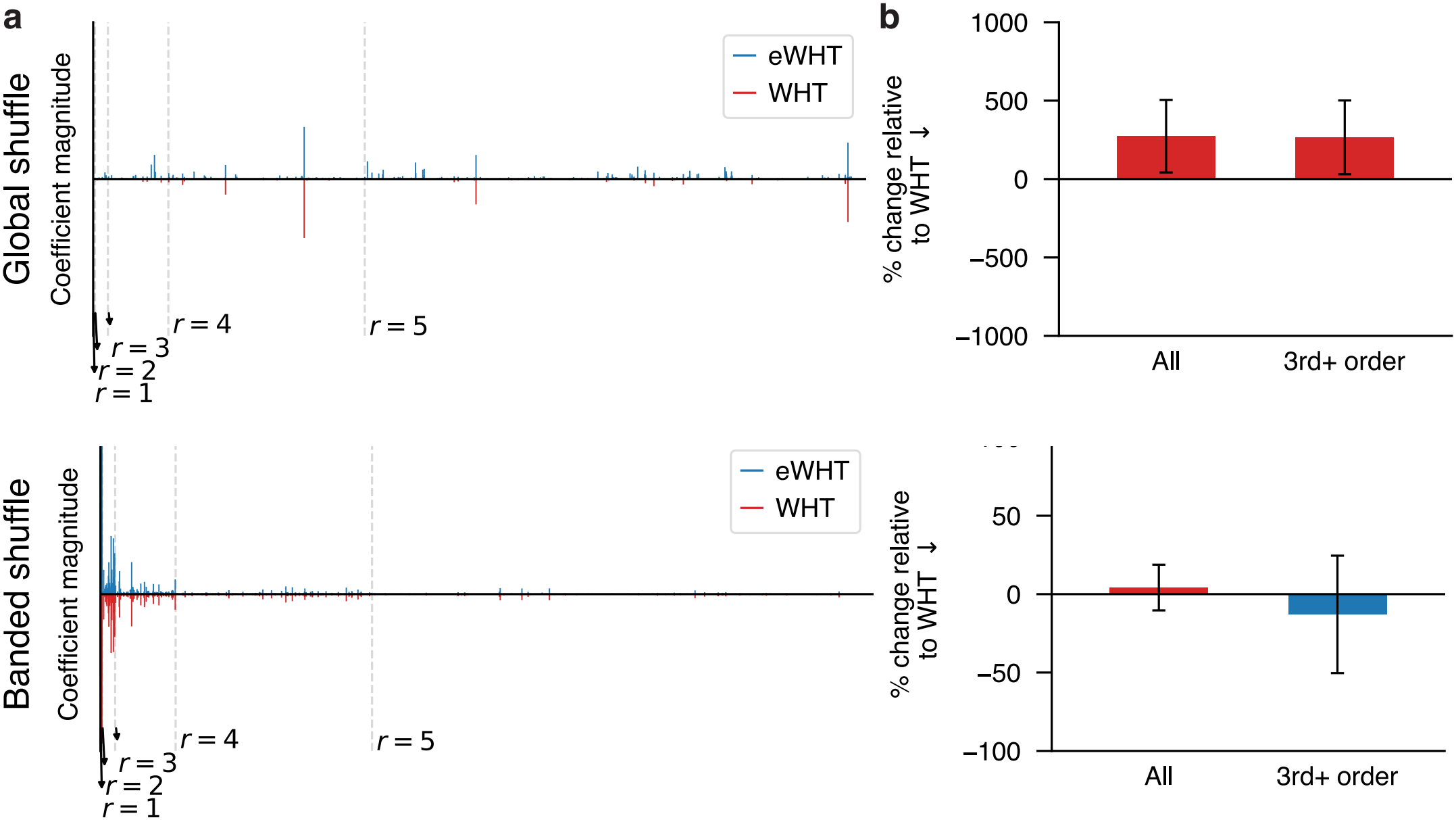
Shuffling the WHT coefficients of the fitness landscape removes or reverses the advantage of eWHT. Comparison of eWHT and WHT on two synthetic landscapes constructed from the experimental fitness landscapes (see Fig. S11 for the experimental landscapes). In the *global shuffle* landscape, the WHT coefficients are randomly shuffled across the entire spectrum and assigned random signs. In the *banded shuffle* landscape, the WHT coefficients are randomly shuffled only within each interaction order and assigned random signs. In both, the evolutionary measure and evaluation sequences are those of the experimental landscape, and only the sequence-to-function mapping differs. Panels **a** and **b** are as in Fig. S11. Relative to the experimental landscapes, the average reduction is reversed under global shuffle and smaller under banded shuffle.

**Figure S13:**
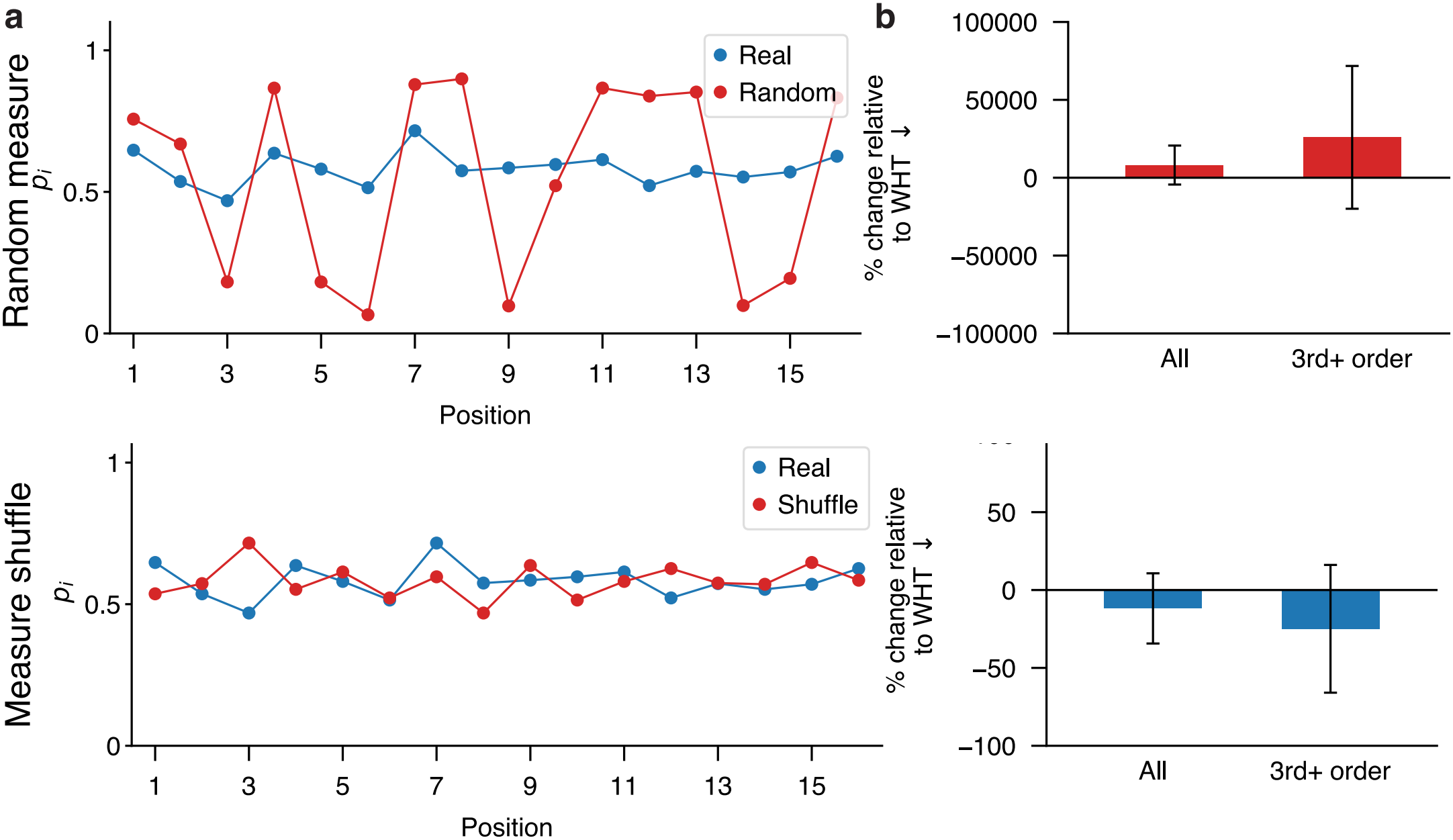
Altering the evolutionary measure weakens or reverses the advantage of eWHT. Comparison of eWHT and WHT on two setups constructed from the experimental fitness landscapes (see Fig. S11 for the experimental landscapes). In both, the fitness landscape and evaluation sequences are those of the experimental landscape, and only the evolutionary measure ***p*** differs. In the *random measure* setup, *p*_*i*_ at each site is drawn uniformly at random from [0.05, 0.95]. In the *measure shuffle* setup, the evolutionary measure ***p*** is randomly shuffled across positions. Panels **a** and **b** are as in Fig. S11. Relative to the experimental landscapes, the average reduction is smaller under measure shuffle and reversed under random measure.

**Table S1:** Language model used to define the evolutionary measure for each landscape, the size of its genotype space, and the absolute Spearman correlation (|*ρ*|) between the score used and experimental fitness. For AntiBERTy, the score is the pseudo-perplexity, and for ESM2-650M, it is the sum of the log-likelihood ratios of the mutations of a variant relative to the wild-type sequence.

| **Landscape** | **Genotype space** | **Model** | **Spearman \|ρ\|** |
|---|---|---|---|
| COV107-23 | 2<sup>10</sup> | AntiBERTy | 0.44 |
| CR9114-H1 | 2<sup>16</sup> | AntiBERTy | 0.34 |
| CR6261-H1 | 2<sup>11</sup> | AntiBERTy | 0.62 |
| CR9114-H3 | 2<sup>16</sup> | AntiBERTy | 0.26 |
| CR6261-H9 | 2<sup>11</sup> | AntiBERTy | 0.53 |
| TagBFP | 2<sup>13</sup> | ESM2 | 0.12 |
| His3p | 2<sup>11</sup> | ESM2 | 0.31 |
| *β*-lactamase | 2<sup>5</sup> | ESM2 | 0.61 |
| Hu-1 | 2<sup>15</sup> | ESM2 | 0.14 |

**Table S2:** Number of coefficients required by WHT and eWHT to reach *R*^2^ = 0.95 on the evolutionarily plausible sequences of each experimental landscape, shown for all coefficients and for the third-order and higher coefficients among them. Positive % reductions indicate that eWHT required fewer coefficients than WHT, while negative % reductions indicate that eWHT required more coefficients than WHT.

|  | **All coefficients** |  |  | **3rd+ order coefficients** |  |  |
|---|---|---|---|---|---|---|
| **Landscape** | **WHT** | **eWHT** | **% reduction** | **WHT** | **eWHT** | **% reduction** |
| COV107-23 | 17 | 12 | 29 | 1 | 0 | 100 |
| CR9114-H1 | 46 | 39 | 15 | 18 | 16 | 11 |
| CR6261-H1 | 567 | 404 | 29 | 515 | 356 | 31 |
| CR9114-H3 | 137 | 133 | 3 | 85 | 78 | 8 |
| CR6261-H9 | 350 | 268 | 23 | 307 | 227 | 26 |
| TagBFP | 26 | 27 | −4 | 15 | 12 | 20 |
| His3p | 374 | 290 | 22 | 321 | 233 | 27 |
| *β*-lactamase | 21 | 7 | 67 | 10 | 1 | 90 |
| Hu-1 | 30 | 28 | 7 | 6 | 5 | 17 |
| **Average** |  |  | 21 ± 21 |  |  | 37 ± 34 |

**Table S3:** Number of coefficients required by WHT and eWHT to reach MSE = 0.05 on high-fitness sequences of each experimental landscape, shown for all coefficients and for the third-order and higher coefficients among them. Positive % reductions indicate that eWHT required fewer coefficients than WHT, while negative % reductions indicate that eWHT required more coefficients than WHT. A dash indicates that WHT required no third-order or higher coefficients, so the reduction is undefined; the average for third-order and higher coefficients is taken over the three landscapes in which it is defined.

|  | **All coefficients** |  |  | **3rd+ order coefficients** |  |  |
|---|---|---|---|---|---|---|
| **Landscape** | **WHT** | **eWHT** | **% reduction** | **WHT** | **eWHT** | **% reduction** |
| COV107-23 | 7 | 7 | 0 | 0 | 0 | — |
| CR9114-H1 | 5 | 4 | 20 | 0 | 0 | — |
| CR6261-H1 | 64 | 37 | 42 | 35 | 15 | 57 |
| CR9114-H3 | 108 | 97 | 10 | 62 | 50 | 19 |
| CR6261-H9 | 226 | 200 | 12 | 187 | 161 | 14 |
| TagBFP | 4 | 5 | −25 | 0 | 0 | — |
| His3p | 5 | 1 | 80 | 0 | 0 | — |
| *β*-lactamase | 1 | 1 | 0 | 0 | 0 | — |
| Hu-1 | 10 | 12 | −20 | 0 | 0 | — |
| **Average** |  |  | 13 ± 32 |  |  | 30 ± 24 |

**Table S4:** Number of coefficients required by WHT and eWHT to reach *R*^2^ = 0.95 on the evolutionarily plausible sequences in the shuffled fitness landscapes, shown for all coefficients and for the third-order and higher coefficients among them. Each synthetic landscape was generated from each fitness landscape using three random seeds.

|  | **All coefficients** |  |  | **3rd+ order coefficients** |  |  |
|---|---|---|---|---|---|---|
| **Landscape** | **WHT** | **eWHT** | **% reduction** | **WHT** | **eWHT** | **% reduction** |
| COV107-23 | 567 ± 9 | 509 ± 8 | 10 ± 0 | 533 ± 4 | 476 ± 7 | 11 ± 1 |
| CR9114-H1 | 36381 ± 157 | 33902 ± 164 | 7 ± 0 | 36301 ± 154 | 33823 ± 152 | 7 ± 0 |
| CR6261-H1 | 1133 ± 8 | 1074 ± 17 | 5 ± 1 | 1094 ± 9 | 1034 ± 18 | 5 ± 1 |
| CR9114-H3 | 36394 ± 149 | 33898 ± 88 | 7 ± 0 | 36313 ± 153 | 33813 ± 85 | 7 ± 0 |
| CR6261-H9 | 1133 ± 30 | 1063 ± 14 | 6 ± 1 | 1093 ± 30 | 1021 ± 13 | 7 ± 1 |
| TagBFP | 4533 ± 63 | 4217 ± 26 | 7 ± 1 | 4482 ± 66 | 4164 ± 21 | 7 ± 1 |
| His3p | 1147 ± 19 | 1057 ± 9 | 8 ± 1 | 1107 ± 17 | 1016 ± 8 | 8 ± 1 |
| *β*-lactamase | 20 ± 4 | 18 ± 4 | 10 ± 2 | 10 ± 2 | 7 ± 3 | 29 ± 20 |
| Hu-1 | 18151 ± 67 | 17048 ± 70 | 6 ± 1 | 18084 ± 76 | 16978 ± 64 | 6 ± 1 |
| **Average** |  |  | 7 ± 2 |  |  | 10 ± 7 |

**Table S5:** Number of coefficients required by WHT and eWHT to reach *R*^2^ = 0.95 on the evolutionarily plausible sequences in the global shuffle landscapes, shown for all coefficients and for the third-order and higher coefficients among them. Each synthetic landscape was generated from each fitness landscape using three random seeds.

|  | **All coefficients** |  |  | **3rd+ order coefficients** |  |  |
|---|---|---|---|---|---|---|
| **Landscape** | **WHT** | **eWHT** | **% reduction** | **WHT** | **eWHT** | **% reduction** |
| COV107-23 | 17 ± 1 | 62 ± 17 | −264 ± 86 | 16 ± 2 | 57 ± 19 | −249 ± 88 |
| CR9114-H1 | 76 ± 1 | 462 ± 89 | −510 ± 114 | 76 ± 1 | 457 ± 93 | −504 ± 120 |
| CR6261-H1 | 228 ± 15 | 358 ± 19 | −57 ± 7 | 221 ± 13 | 339 ± 15 | −53 ± 7 |
| CR9114-H3 | 143 ± 0 | 830 ± 52 | −481 ± 36 | 143 ± 0 | 825 ± 54 | −477 ± 38 |
| CR6261-H9 | 213 ± 3 | 367 ± 29 | −72 ± 12 | 206 ± 6 | 347 ± 24 | −68 ± 8 |
| TagBFP | 57 ± 3 | 295 ± 47 | −415 ± 78 | 56 ± 2 | 291 ± 46 | −416 ± 81 |
| His3p | 271 ± 28 | 415 ± 40 | −53 ± 13 | 263 ± 28 | 389 ± 43 | −48 ± 13 |
| *β*-lactamase | 14 ± 1 | 15 ± 3 | −10 ± 25 | 8 ± 1 | 7 ± 3 | 18 ± 28 |
| Hu-1 | 28 ± 1 | 193 ± 22 | −597 ± 72 | 28 ± 1 | 192 ± 23 | −594 ± 76 |
| **Average** |  |  | −273 ± 231 |  |  | −266 ± 235 |

**Table S6:**
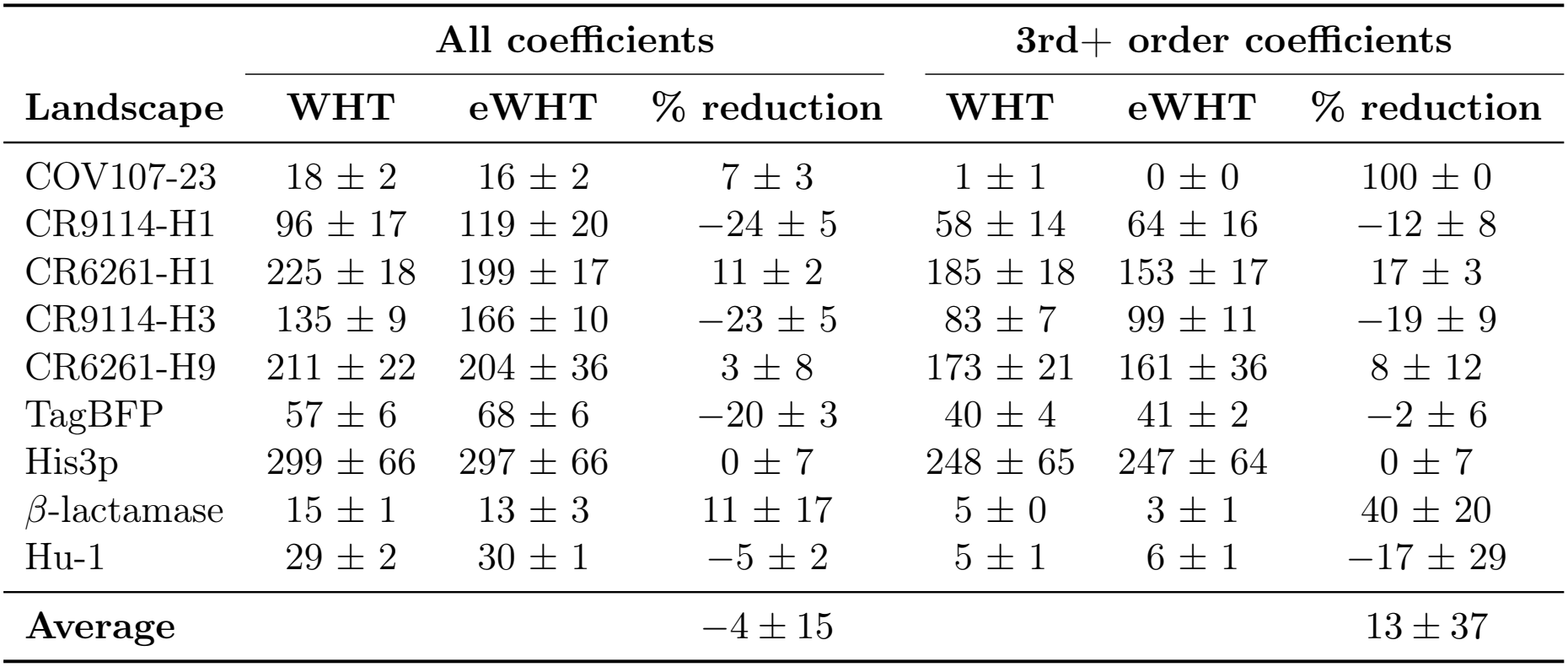
Number of coefficients required by WHT and eWHT to reach *R*^2^ = 0.95 on the evolutionarily plausible sequences in the banded shuffle landscapes, shown for all coefficients and for the third-order and higher coefficients among them. Each synthetic landscape was generated from each fitness landscape using three random seeds.

**Table S7:**
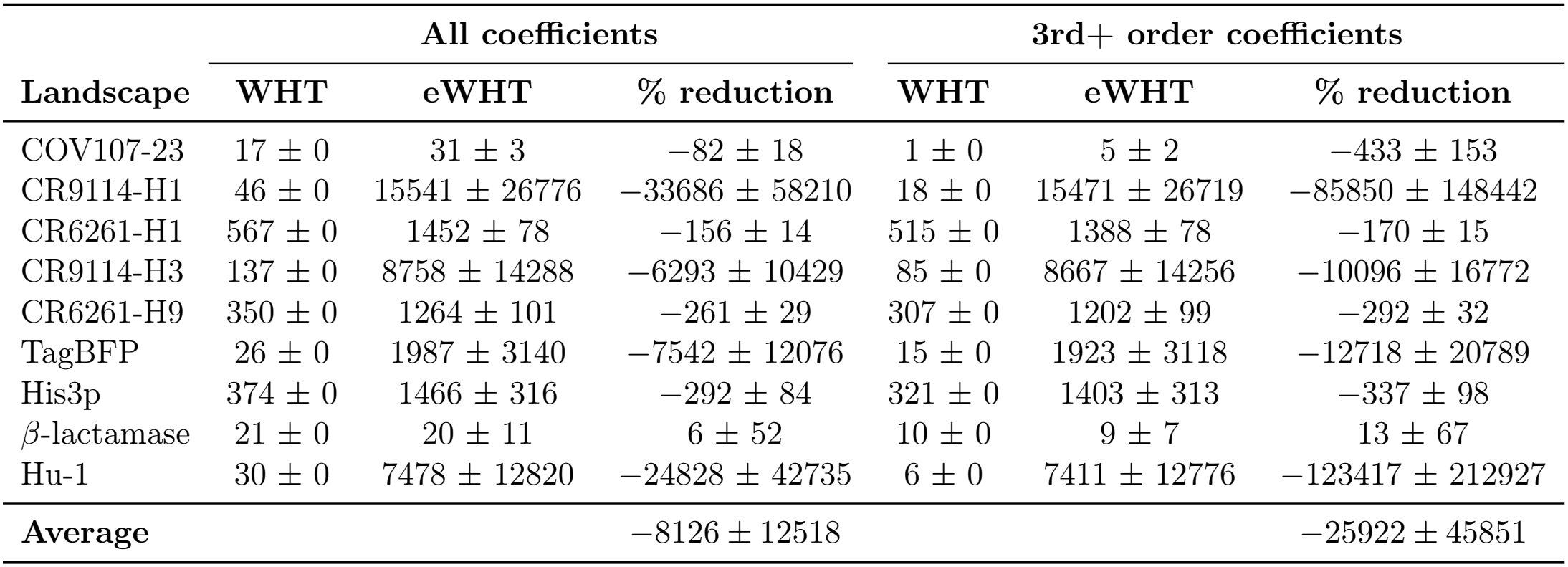
Number of coefficients required by WHT and eWHT to reach *R*^2^ = 0.95 on the evolutionarily plausible sequences in the random measure setup, shown for all coefficients and for the third-order and higher coefficients among them. Each random measure was generated for each fitness landscape using three random seeds. Since the random measure affects only ***p***, the WHT counts are the same as in the real fitness landscape setup.

**Table S8:** Number of coefficients required by WHT and eWHT to reach *R*^2^ = 0.95 on the evolutionarily plausible sequences in the measure shuffle setup, shown for all coefficients and for the third-order and higher coefficients among them. Each measure shuffle was generated for each fitness landscape using three random seeds. Since the measure shuffle affects only ***p***, the WHT counts are the same as in the real fitness landscape setup.

|  | **All coefficients** |  |  | **3rd+ order coefficients** |  |  |
|---|---|---|---|---|---|---|
| **Landscape** | **WHT** | **eWHT** | **% reduction** | **WHT** | **eWHT** | **% reduction** |
| COV107-23 | 17 ± 0 | 13 ± 1 | 25 ± 3 | 1 ± 0 | 0 ± 0 | 100 ± 0 |
| CR9114-H1 | 46 ± 0 | 44 ± 3 | 4 ± 6 | 18 ± 0 | 18 ± 0 | 0 ± 0 |
| CR6261-H1 | 567 ± 0 | 440 ± 8 | 22 ± 1 | 515 ± 0 | 388 ± 8 | 25 ± 2 |
| CR9114-H3 | 137 ± 0 | 149 ± 4 | −9 ± 3 | 85 ± 0 | 91 ± 3 | −7 ± 3 |
| CR6261-H9 | 350 ± 0 | 303 ± 19 | 13 ± 6 | 307 ± 0 | 260 ± 18 | 15 ± 6 |
| TagBFP | 26 ± 0 | 27 ± 0 | −4 ± 0 | 15 ± 0 | 12 ± 0 | 20 ± 0 |
| His3p | 374 ± 0 | 425 ± 52 | −14 ± 14 | 321 ± 0 | 370 ± 53 | −15 ± 16 |
| *β*-lactamase | 21 ± 0 | 8 ± 3 | 60 ± 15 | 10 ± 0 | 1 ± 2 | 87 ± 15 |
| Hu-1 | 30 ± 0 | 28 ± 0 | 7 ± 0 | 6 ± 0 | 6 ± 0 | 0 ± 0 |
| **Average** |  |  | 12 ± 23 |  |  | 25 ± 41 |

## Notes

https://github.com/amirgroup-codes/eWHT_expts/

https://github.com/amirgroup-codes/ewht

https://figshare.com/articles/dataset/eWHT_Data/32869433

### Competing Interest Statement

The authors have declared no competing interest.

### Summary of Updates

Paper updated in accordance with new results

## References

[1] Daniel M. Weinreich, Nigel F. Delaney, Mark A. DePristo, and Daniel L. Hartl. Darwinian evolution can follow only very few mutational paths to fitter proteins. Science, 312(5770):111–114, 2006.

[2] Patrick C Phillips. Epistasis—the essential role of gene interactions in the structure and evolution of genetic systems. Nature Reviews Genetics, 9(11):855–867, 2008.

[3] Frank J. Poelwijk, Michael Socolich, and Rama Ranganathan. Learning the pattern of epistasis linking genotype and phenotype in a protein. Nature Communications, 10(1), 2019.

[4] Douglas M Fowler and Stanley Fields. Deep mutational scanning: a new style of protein science. Nature Methods, 11(8):801–807, 2014.

[5] Douglas M Fowler, Jason J Stephany, and Stanley Fields. Measuring the activity of protein variants on a large scale using deep mutational scanning. Nature Protocols, 9(9):2267–2284, 2014.

[6] Allison Judge, Banumathi Sankaran, Liya Hu, Murugesan Palaniappan, André Birgy, BV Venkataram Prasad, and Timothy Palzkill. Network of epistatic interactions in an enzyme active site revealed by large-scale deep mutational scanning. Proceedings of the National Academy of Sciences, 121(12):e2313513121, 2024.

[7] Daniel M Weinreich, Yinghong Lan, C Scott Wylie, and Robert B. Heckendorn. Should evolutionary geneticists worry about higher-order epistasis? Current Opinion in Genetics & Development, 23(6):700–707, 2013.

[8] Frank J Poelwijk, Vinod Krishna, and Rama Ranganathan. The context-dependence of mutations: a linkage of formalisms. PLoS Computational Biology, 12(6):e1004771, 2016.

[9] Alief Moulana, Thomas Dupic, Angela M. Phillips, Jeffrey Chang, Serafina Nieves, Anne A. Roffler, Allison J. Greaney, Tyler N. Starr, Jesse D. Bloom, and Michael M. Desai. Compensatory epistasis maintains ACE2 affinity in SARS-CoV-2 Omicron BA.1. Nature Communications, 13(1), 2022.

[10] Steven Schulz, Timothy J. C. Tan, Nicholas C. Wu, and Shenshen Wang. Epistatic hotspots organize antibody fitness landscape and boost evolvability. Proceedings of the National Academy of Sciences, 122(2), 2025.

[11] Thomas Dupic, Angela M. Phillips, and Michael M. Desai. Protein sequence landscapes are not so simple: on reference-free versus reference-based inference. bioRxiv, 2024.

[12] Yeonwoo Park, Brian P. H. Metzger, and Joseph W. Thornton. The simplicity of protein sequence-function relationships. Nature Communications, 15(1), 2024.

[13] Karen S. Sarkisyan, Dmitry A. Bolotin, Margarita V. Meer, Dinara R. Usmanova, Alexander S. Mishin, George V. Sharonov, Dmitry N. Ivankov, Nina G. Bozhanova, Mikhail S. Baranov, Onuralp Soylemez, Natalya S. Bogatyreva, Peter K. Vlasov, Evgeny S. Egorov, Maria D. Logacheva, Alexey S. Kondrashov, Dmitry M. Chudakov, Ekaterina V. Putintseva, Ilgar Z. Mamedov, Dan S. Tawfik, Konstantin A. Lukyanov, and Fyodor A. Kondrashov. Local fitness landscape of the green fluorescent protein. Nature, 533(7603):397–401, 2016.

[14] Nicholas C Wu, Lei Dai, C Anders Olson, James O Lloyd-Smith, and Ren Sun. Adaptation in protein fitness landscapes is facilitated by indirect paths. eLife, 5, 2016.

[15] David H Brookes, Amirali Aghazadeh, and Jennifer Listgarten. On the sparsity of fitness functions and implications for learning. Proceedings of the National Academy of Sciences, 119(1):e2109649118, 2022.

[16] Jakub Otwinowski, David M McCandlish, and Joshua B Plotkin. Inferring the shape of global epistasis. Proceedings of the National Academy of Sciences, 115(32):E7550–E7558, 2018.

[17] Amirali Aghazadeh, Hunter Nisonoff, Orhan Ocal, David H Brookes, Yijie Huang, O Ozan Koyluoglu, Jennifer Listgarten, and Kannan Ramchandran. Epistatic net allows the sparse spectral regularization of deep neural networks for inferring fitness functions. Nature Communications, 12(1):5225, 2021.

[18] Darin Tsui, Aryan Musharaf, Yigit E. Erginbas, Justin S. Kang, and Amirali Aghazadeh. SHAP zero explains biological sequence models with near-zero marginal cost for future queries. In Advances in Neural Information Processing Systems, volume 38, pages 83840–83891, 2025.

[19] Justin S. Kang, Darin Tsui, Yigit Efe Erginbas, Landon Butler, Amirali Aghazadeh, and Kannan Ramchandran. Spectral sparsity: A unifying framework for scalable model interpretability using codes. IEEE BITS the Information Theory Magazine, page 1–14, 2026.

[20] Darin Tsui and Amirali Aghazadeh. On recovering higher-order interactions from protein language models. In ICLR 2024 Workshop on Generative and Experimental Perspectives for Biomolecular Design, 2024.

[21] Victoria O. Pokusaeva, Dinara R. Usmanova, Ekaterina V. Putintseva, Lorena Espinar, Karen S. Sarkisyan, Alexander S. Mishin, Natalya S. Bogatyreva, Dmitry N. Ivankov, Arseniy V. Akopyan, Sergey Ya. Avvakumov, Inna S. Povolotskaya, Guillaume J. Filion, Lucas B. Carey, and Fyodor A. Kondrashov. An experimental assay of the interactions of amino acids from orthologous sequences shaping a complex fitness landscape. PLOS Genetics, 15(4):e1008079, 2019.

[22] Gabriel J. Rocklin, Tamuka M. Chidyausiku, Inna Goreshnik, Alex Ford, Scott Houliston, Alexander Lemak, Lauren Carter, Rashmi Ravichandran, Vikram K. Mulligan, Aaron Chevalier, Cheryl H. Arrowsmith, and David Baker. Global analysis of protein folding using massively parallel design, synthesis, and testing. Science, 357(6347):168–175, 2017.

[23] C Clark Cockerham. An extension of the concept of partitioning hereditary variance for analysis of covariances among relatives when epistasis is present. Genetics, 39(6):859–882, 1954.

[24] José M Alvarez-Castro and Orjan Carlborg. A unified model for functional and statistical epistasis and its application in quantitative trait loci analysis. Genetics, 176(2):1151–1167, 2007.

[25] Peter F. Stadler, Rudi Seitz, and Günter P. Wagner. Population dependent fourier decomposition of fitness landscapes over recombination spaces: Evolvability of complex characters. Bulletin of Mathematical Biology, 62(3):399–428, 2000.

[26] Jeffrey A. Ruffolo, Jeffrey J. Gray, and Jeremias Sulam. Deciphering antibody affinity maturation with language models and weakly supervised learning. arXiv preprint arXiv:2112.07782, 2021.

[27] Angela M Phillips, Katherine R Lawrence, Alief Moulana, Thomas Dupic, Jeffrey Chang, Milo S Johnson, Ivana Cvijovic, Thierry Mora, Aleksandra M Walczak, and Michael M Desai. Binding affinity landscapes constrain the evolution of broadly neutralizing anti-influenza antibodies. eLife, 10, 2021.

[28] Zeming Lin, Halil Akin, Roshan Rao, Brian Hie, Zhongkai Zhu, Wenting Lu, Nikita Smetanin, Robert Verkuil, Ori Kabeli, Yaniv Shmueli, et al. Evolutionary-scale prediction of atomic-level protein structure with a language model. Science, 379(6637):1123–1130, 2023.

[29] Leland McInnes, John Healy, Nathaniel Saul, and Lukas Großberger. UMAP: Uniform manifold approximation and projection. Journal of Open Source Software, 3(29):861, 2018.

[30] Michael Chungyoun, Jeffrey Ruffolo, and Jeffrey Gray. FLAb: Benchmarking deep learning methods for antibody fitness prediction. In NeurIPS 2023 Workshop on Machine Learning in Structural Biology, 2023.

[31] Michael Chungyoun and Jeffrey Gray. Fitness landscape for antibodies 2: Benchmarking reveals that protein AI models cannot yet consistently predict developability properties. bioRxiv, 2025.

[32] Joshua Meier, Roshan Rao, Robert Verkuil, Jason Liu, Tom Sercu, and Alex Rives. Language models enable zero-shot prediction of the effects of mutations on protein function. In Advances in Neural Information Processing Systems, volume 34, pages 29287–29303, 2021.

[33] Pascal Notin, Aaron Kollasch, Daniel Ritter, Lood van Niekerk, Steffanie Paul, Han Spinner, Nathan Rollins, Ada Shaw, Rose Orenbuch, Ruben Weitzman, Jonathan Frazer, Mafalda Dias, Dinko Franceschi, Yarin Gal, and Debora Marks. ProteinGym: Large-scale benchmarks for protein fitness prediction and design. In Advances in Neural Information Processing Systems, volume 36, pages 64331–64379, 2023.

[34] Robert Tibshirani. Regression shrinkage and selection via the lasso. Journal of the Royal Statistical Society Series B: Statistical Methodology, 58(1):267–288, 1996.

[35] Timothy J. C. Tan, Meng Yuan, Kaylee Kuzelka, Gilberto C. Padron, Jacob R. Beal, Xin Chen, Yiquan Wang, Joel Rivera-Cardona, Xueyong Zhu, Beth M. Stadtmueller, Christopher B. Brooke, Ian A. Wilson, and Nicholas C. Wu. Sequence signatures of two public antibody clonotypes that bind SARS-CoV-2 receptor binding domain. Nature Communications, 12(1), 2021.

[36] Cyrille Dreyfus, Nick S. Laursen, Ted Kwaks, David Zuijdgeest, Reza Khayat, Damian C. Ekiert, Jeong Hyun Lee, Zoltan Metlagel, Miriam V. Bujny, Mandy Jongeneelen, Remko van der Vlugt, Mohammed Lamrani, Hans J. W. M. Korse, Eric Geelen, Özcan Sahin, Martijn Sieuwerts, Just P. J. Brakenhoff, Ronald Vogels, Olive T. W. Li, Leo L. M. Poon, Malik Peiris, Wouter Koudstaal, Andrew B. Ward, Ian A. Wilson, Jaap Goudsmit, and Robert H. E. Friesen. Highly conserved protective epitopes on influenza B viruses. Science, 337(6100):1343–1348, 2012.

[37] Damian C. Ekiert, Gira Bhabha, Marc-Andre Elsliger, Robert H. E. Friesen, Mandy Jongeneelen, Mark Throsby, Jaap Goudsmit, and Ian A. Wilson. Antibody recognition of a highly conserved influenza virus epitope. Science, 324(5924):246–251, 2009.

[38] Christian Dallago, Jody Mou, Kadina E Johnston, Bruce J Wittmann, Nicholas Bhattacharya, Samuel Goldman, Ali Madani, and Kevin K Yang. FLIP: Benchmark tasks in fitness landscape inference for proteins. NeurIPS Datasets Benchmarks, 2021.

[39] Zachary R Sailer and Michael J Harms. Detecting high-order epistasis in nonlinear genotype-phenotype maps. Genetics, 205(3):1079–1088, 2017.

[40] Thomas A Hopf, Charlotta Schärfe, João Rodrigues, Anna G Green, Oliver Kohlbacher, Chris Sander, Alexandre M J J Bonvin, and Debora S Marks. Sequence co-evolution gives 3D contacts and structures of protein complexes. eLife, 3, 2014.

[41] Thomas A Hopf, John B Ingraham, Frank J Poelwijk, Charlotta PI Schärfe, Michael Springer, Chris Sander, and Debora S Marks. Mutation effects predicted from sequence co-variation. Nature Biotechnology, 35(2):128–135, 2017.

[42] Darin Tsui, Kunal Talreja, and Amirali Aghazadeh. Efficient algorithm for sparse Fourier transform of generalized q-ary functions. In 2025 IEEE Information Theory Workshop (ITW), page 1–6. IEEE, 2025.

[43] Ryan O’Donnell. Analysis of Boolean Functions. Cambridge University Press, Cambridge, 2014.

[44] Emmanuel Candes, Justin Romberg, and Terence Tao. Robust uncertainty principles: Exact signal reconstruction from highly incomplete frequency information, 2004.

